# Hypomagnetic field inhibits seed germination via ROS signaling and reprogramming transcriptome in *Arabidopsis thaliana*

**DOI:** 10.64898/2026.09.29.755266

**Authors:** Yang Gu, Wenjuan Wu, Xianyang Su, Yanwen Fang, Yidong Shen, Jirong Huang, Xiang Xu

## Abstract

The geomagnetic field (GMF) is a ubiquitous yet poorly understood environmental factor that affects plant growth and development. It has been suggested that the blue-light photoreceptor cryptochrome (CRY) functions as a critical magnetic-field sensor in birds and insects through the radical-pair mechanism (RPM). However, whether CRY acts as a *bona fide* transducer of GMF signals and what are the downstream cascades remain largely elusive in plants. To address these key issues, we generated hypomagnetic field (HMF) via Helmholtz coils and passive magnetic shielding to investigate the GMF deprivation effects in *Arabidopsis thaliana*. HMF significantly delays the germination speed without altering the final germination rate, suggesting that HMF specifically modulates germination kinetics rather than seed viability. Time-resolved transcriptomic profiling revealed a coordinated transcriptional shift characterized by downregulation of growth-promoting genes and concurrent upregulation of defense-related genes under HMF. This biphasic transcriptional reprogramming coincides with a significant increase of reactive oxygen species (ROS), linking redox perturbation to the germination delay. Consistently, supplementation with ROS-scavenging antioxidants (e.g., reduced glutathione and ascorbic acid) rescues both germination speed and the misregulated expression of a subset of HMF-responsive genes, suggesting the central role of ROS in mediating HMF bioeffects. Genetic analysis using *cry1 cry2* double mutants further indicated that HMF-induced germination delay operates through both CRY-dependent and -independent signaling pathways, suggesting the involvement of additional magnetosensitive modules beyond the canonical CRY-based RPM. These findings suggest that the GMF acts as a positive environmental cue that fine tunes the growth-defense trade-off through a redox-dependent signaling.

## Introduction

The geomagnetic field (GMF), a pervasive yet often overlooked important environmental factor on the Earth, is crucial not only for shielding life from the solar wind and cosmic radiation but also for influencing animal migration and regulating fundamental biological processes, such as cell proliferation, differentiation, and circadian rhythms in living organisms (Erdmann *et al*., 2021; Lammer *et al*., 2018; Sarimov *et al*., 2023). One of the most efficient approaches to evaluate the GMF role in living organisms is to study their phenotypes and changes at the molecular level in a near-zero magnetic field (NZMF) or hypomagnetic field (HMF) environment, in which the most commonly used threshold is below 300 nT (Sarimov *et al*., 2023). To date, it has been well demonstrated that HMF exposure has multiple negative impacts on animal organ systems including reproductive, central nervous, cardiovascular and musculoskeletal systems (Tian *et al*., 2024). These findings have laid a strong foundation to systematically assess the potential health risks for humans during future deep-space exploration. In contrast, biological effects of HMF on growth and development of higher plants remain largely underexplored. This knowledge gap is especially significant given the growing interest in cultivating plants in extraterrestrial environments, where magnetic fields are absent or significantly altered.

When plants are subjected to grow under HMF conditions, a range of phenotypic changes have been reported, including elongated hypocotyls, reduced overall growth, and delayed flowering (Lebedev S I, 1977; Maffei, 2014; Xu *et al*., 2012). However, experimental results in HMF are sometimes controversial or difficult to reproduce. These discrepancies are probably attributable to variables such as plant species, genotype, treatment duration, and growth conditions (Harris *et al*., 2009; Rakosy-Tican *et al*., 2005). For example, HMF treatment has been shown to increase, decrease and have no significant effect on germination rate in soybean, *Arabidopsis*, and sunflower, respectively (Belyavskaya, 2004; Dhiman *et al*., 2023; Fischer *et al*., 2004; Mo *et al*., 2011). In contrast, exposure to moderate static magnetic field (SMF) generally promotes seed germination across various species, such as quinoa, sunflower, maize, and rice by enhancing water uptake, enzymatic activity, and metabolic rates (Luo *et al*., 2022; Vashisth and Joshi, 2017; Vashisth and Nagarajan, 2010; S. Wang *et al*., 2024).

At the molecular level, reactive oxygen species (ROS) are hypothesized to be key signaling intermediates in magnetic field responses (Li *et al*., 2025; Paponov *et al*., 2021; Pszczółkowski *et al*., 2023; Sarraf *et al*., 2020; Xu *et al*., 2026). ROS can be intricately generated through the cryptochrome-dependent radical pair mechanism (RPM), which is the most established quantum-chemical model of how eukaryotes sense magnetic field. In plants, cryptochromes form photoinduced radical pairs upon blue-light absorption; at the subsequent reoxidation step, the flavin semiquinone can interact with molecular oxygen, generating superoxide (Maffei, 2025). In addition, under hypomagnetic conditions ROS can be produced through multiple pathways, such as mitochondrial respiration, peroxisomes, and NADPH oxidase in the plasma membrane (Maffei, 2025). In *Arabidopsis* seedlings, HMF exposure led to a lower level of H_2_O_2_ than GMF, alongside reduced expression of ROS-related genes and lower antioxidant polyphenol content (Parmagnani *et al*., 2022). This aligns with the established role of optimal ROS concentration window in promoting seed dormancy release and germination (Y. Wang *et al*., 2025; Xie *et al*., 2025). However, the precise mechanistic link between HMF exposure and ROS dynamics especially during seed germination process remains poorly defined.

In this study, we investigated the effect of HMF exposure on seed germination in *Arabidopsis thaliana*. We found that HMF significantly delays germination. Time-course transcriptomic analysis revealed a limited number of differentially expressed genes (DEGs), many of which are associated with stress responses. Notably, HMF induced elevated accumulation of ROS. This germination delay was partially rescued by treatment with ROS scavengers. Collectively, these findings suggest that ROS accumulation plays a key role in mediating the inhibitory effects of HMF on early seedling development, providing new insights into plant adaptation to altered magnetic environments.

## Results

### HMF exposure delays growth and development throughout the plant life cycle

To investigate the effects of HMF conditions on plant growth and development, we established an HMF environment using a triaxial Helmholtz coil system to actively compensate the local GMF. An identical coil system without current input was used as a matched GMF control. Both systems were placed in the same growth chamber and were employed reciprocally for the HMF and control treatments. Samples were positioned at the center of the coils within a 10 cm side-length cubic space (Supplementary Fig. 1). The field intensity of this space was kept below 0.2 μT when the central point field was near zero. Continuous recording of the central point field indicated periodic variation in the local GMF (Supplementary Fig. 2A-D). Correspondingly, in the HMF, the field components in the x, y and z directions were kept around zero, with the total field remaining at approximately 200 nT during daytime (Supplementary Fig. 2E-H).

Our results showed that HMF exposure induced multiple phenotypic changes in *Arabidopsis* plants, compared to the control. Time-course imaging and quantification revealed a significantly lower germination rates between 44 and 48 hours post-sowing in HMF than in GMF (Fig. 1A, B, Supplementary Fig. 3), indicating that magnetic field deprivation impedes this early developmental transition. Nevertheless, the final germination rate was comparable between the two conditions. In addition, germination assays of other four ecotypes yielded similar results to those of Col-0 with the except of Ca-0 (Supplementary Fig. 4). These results suggest that HMF-induced inhibition of seed germination is under genetic control. To further validate the delay in seed germination under HMF, we carried out experiments using a passive shielding (μ-metal shielding) device, and obtained results identical to those observed in coils (Supplementary Fig. 5A, B). Furthermore, the delayed germination induced by HMF was completely reversed by introducing a magnet to recover the field intensity to GMF (Supplementary Fig. 5C, D). Finally, we investigated the effect of 600 mT magnetic field on seed germination. Our data showed that seed germination was significantly enhanced at various time points (Supplementary Fig. 6A). Taken together, these results suggest that seed germination is sensitive to external magnetic fields.

**Fig. 1.**
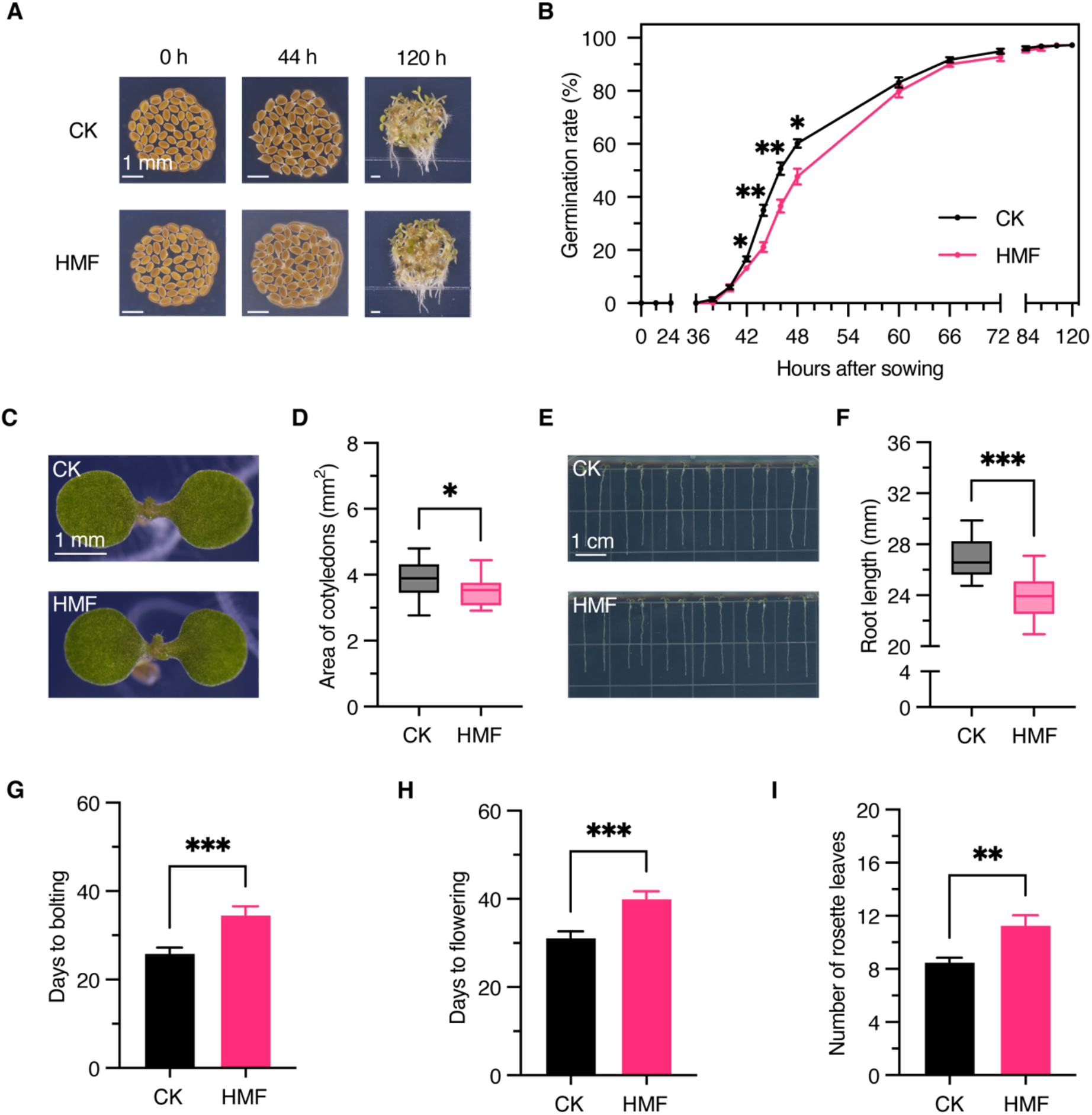
Effects of HMF on *Arabidopsis* seed germination and seedling growth. (A, B) Images (A) and quantification (B) of seed germination at various indicated times. Data are shown means ± SE (n = 4). Stars indicate significant differences between GMF (CK) and HMF (Student’s *t*-test). *, *p* < 0.05; **, *p* < 0.01. Scale bar, 1 mm. (C, D) Cotyledon size (C) and quantification (D) of 7-day-old seedlings grown under control and HMF conditions. Data are presented mean ± SE (n ≥ 36). *, significant difference between CK and HMF (Student’s *t*-test, *p* < 0.05). Scale bar, 1 mm. (E, F) Images (E) and primary root length (F) of 7-day-old seedlings. Data are presented mean ± SE (n ≥ 24). ***, significant difference between CK and HMF (Student’s *t*-test, *p* < 0.001). Scale bar, 1 cm. (G-I) Effects of HMF on bolting (G) and flowering (H, I). Flowering times were analyzed by days (H) and rosette leaves (I) from sowing to flowering. Data are presented as mean ± SE (n ≥ 24). Statistical analysis was performed using Student’s *t*-test (**, *p* < 0.01; ***, *p* < 0.001).

In 7-day-old seedlings, cotyledon size was visibly smaller in HMF than in GMF (Fig. 1C), and quantitative analysis confirmed a significant decrease in cotyledon area (Fig. 1D), suggesting impaired post-germination growth. In addition, HMF-grown seedlings exhibited a pronounced reduction in primary root length, compared to GMF-grown controls (Fig. 1E, F), implying that root meristem activity or elongation is sensitive to magnetic field deprivation.

Besides early developmental defects, HMF exposure also led to a delayed transition from vegetative to reproductive growth. Plants grown under HMF exhibited markedly later bolting and flowering times than those under GMF (Fig. 1G, H). The delayed flowering was further underscored by a significant increase in the number of rosette leaves produced prior to flowering (Fig. 1I). Together, these findings indicate that HMF broadly inhibits *Arabidopsis* growth and development.

### HMF-induced transcriptomic changes during seed germination

To explore the molecular basis of HMF-induced phenotypic changes in *Arabidopsis*, we performed time-course transcriptomic profiling of germinating seeds under GMF and HMF conditions. As illustrated in Fig. 2A, seeds were stratified at 4 °C for 48 hours, then transferred to either GMF or HMF environments. Samples were collected at 28 h, 36 h, and 44 h after sowing to capture key stages of early germination. RNA-Seq data analysis was summarized in Supplementary Table 1. Principal component analysis (PCA) revealed that samples were clearly separated by the germination time, and clustered more closely by the same magnetic field condition at each germination time (Fig. 2B). Consistently, analysis of the sample distance heatmap showed high similarity between HMF and GMF samples at the same time point and among biological replicates (Supplementary Fig. 7). Taken together, these results suggest that transcriptional profiles were regulated predominantly by the seed germination stage while finely by the magnetic field treatment.

**Fig. 2.**
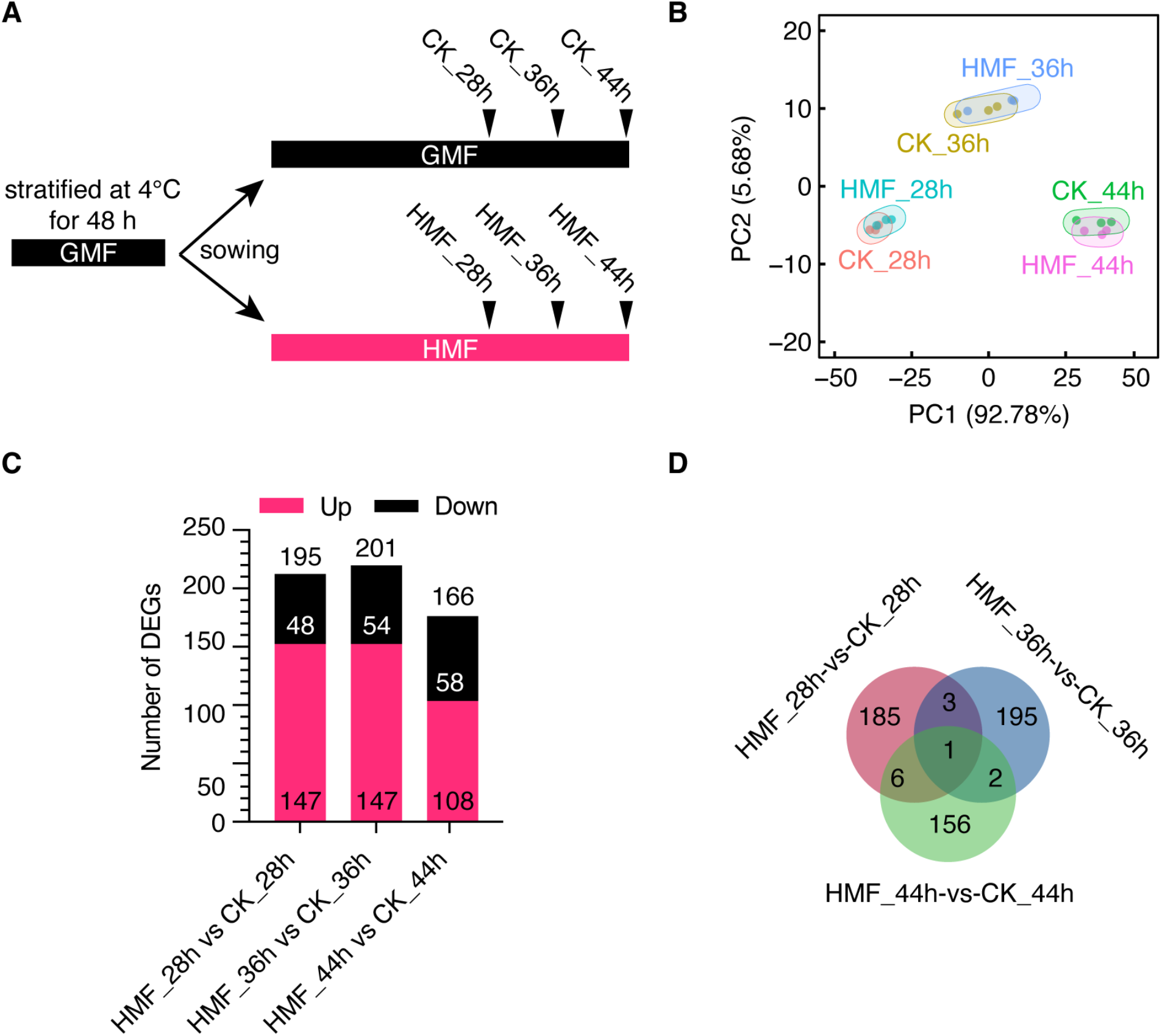
Transcriptomic analysis of germinating seeds in GMF and HMF. (A) Schematic diagram of experimental procedure and sampling times. Seeds stratified at 4°C for 48 hours were germinated under HMF and GMF. Samples were collected at 28 h, 36 h, and 44 h after sowing. (B) Principal component analysis (PCA) of transcriptomic variation among different treatments at three germination points. Each treatment was repeated three times. (C) Differentially expressed genes (DEGs) between HMF and GMF at each sampling time. (D) Venn diagram showing the overlap of DEGs between HMF and CK at 28 h, 36 h, and 44 h after sowing.

DESeq2 analysis was employed to calculate gene expression levels in the HMF-treated seeds relative to the corresponding control seeds (Supplementary Table 2). A total of 195, 201, and 166 differentially expressed genes (DEGs) with fold change ≥1.5 (p < 0.05) in HMF samples were detected at 28 h, 36 h, and 44 h after sowing, respectively, and majority of these DEGs were upregulated under HMF (Fig. 2C, Fig. 3A-C). The heatmap of the top ten DEGs reveals no overlap at each time point (Fig. 3D-F), indicating that HMF-regulated transcriptional profiles are largely affected by seed germination stages. However, among these top DEGs, we found several key genes involved in stress responses during germination. For instance, at 28 h, *PHOSPHATASE 2C5* (*PP2C5*) and *HVA22 HOMOLOGUE E* (*HVA22E*) were upregulated, while *WRKY41* was downregulated (Chen *et al*., 2002; Diao *et al*., 2024; Wang *et al*., 2023). By 36 h, *DIRIGENT PROTEIN 25* (*DIR25*) showed significant upregulation (Fonseca *et al*., 2025). At 44 h, *THAUMATIN-LIKE PROTEIN 3* (*TLP-3*) and *REDOX RESPONSIVE TRANSCRIPTION FACTOR 1* (*RRTF1*) were upregulated (Hu and Reddy, 1997; W. Zhou *et al*., 2019) (Fig. 3D-F). To validate the reliability of the transcriptomic analysis, a subset of DEGs was randomly selected for qRT-PCR verification. The expression levels detected by qRT-PCR were consistent with the RNA-seq data, confirming the reliability of the sequencing analysis (Supplementary Fig. 8).

**Fig. 3.**
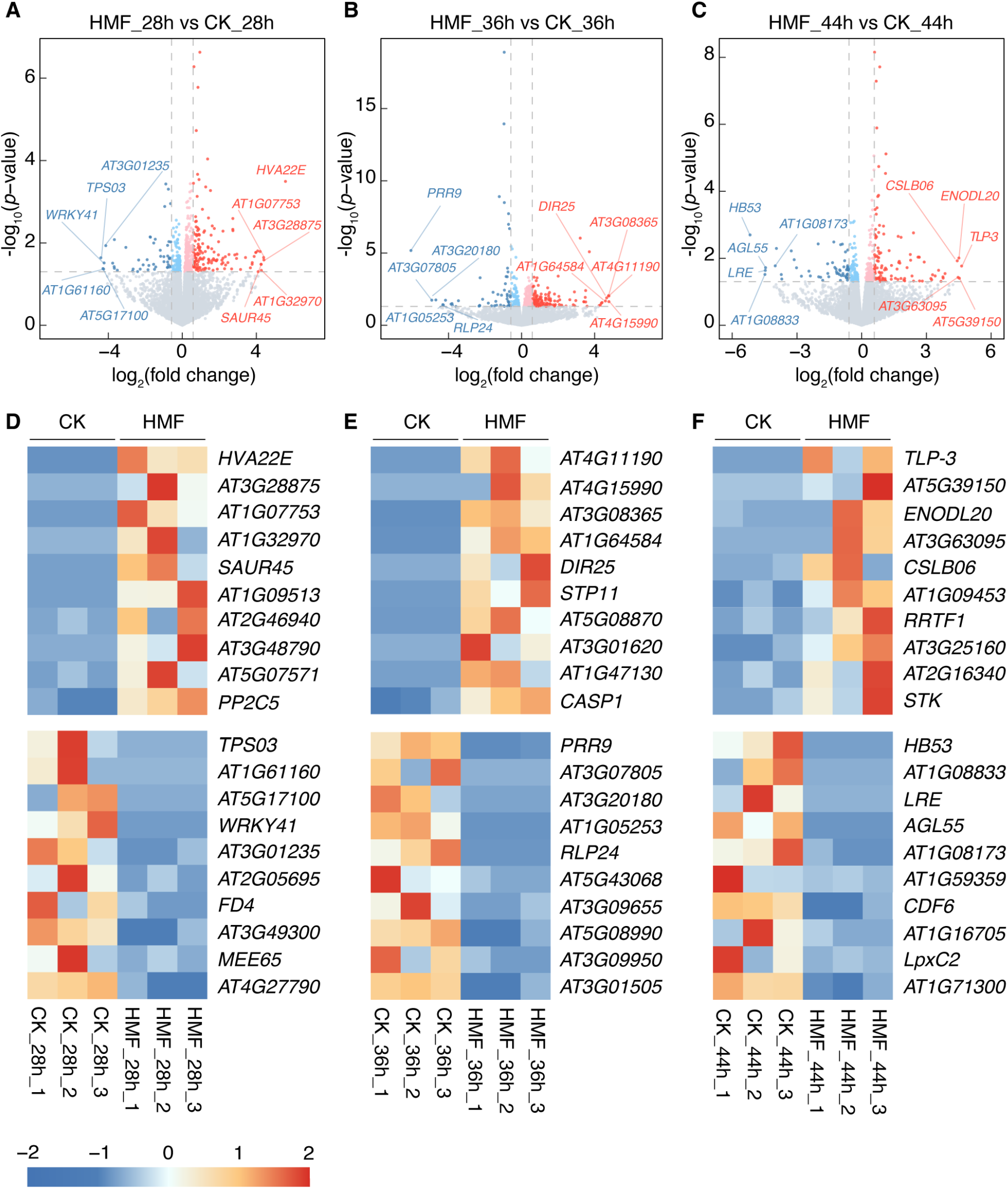
Analysis of DEGs up-or down-regulated by HMF at 28 h, 36 h, and 44 h after sowing. (A-C) Volcano plots of DEGs at 28 h (A), 36 h (B), and 44 h (C) of HMF treatment. Each dot represents a single gene. Red and blue dots indicate significantly up- and down-regulated DEGs, respectively, with fold changes larger than 1.5. Translucent colors indicate fold changes less than 1.5. Gray dots represent non-significant changes. Top 5 DEGs are labeled with gene names or loci. (D-F) Heatmaps show the top 10 DEGs at 28 h (D), 36 h (E), and 44 h (F) of HMF treatment. Red and blue indicate up- and downregulated, respectively. Genes are clustered by expression patterns, and color represents normalized expression values (Z-scores).

Further analysis using a Venn diagram showed limited overlap of DEGs across the three time points. We found that only one DEG was common to all time points, while three were shared between 28 h and 36 h, two between 36 h and 44 h, and six between 28 h and 44 h (Fig. 2D). Notably, the majority of these shared genes remain functionally uncharacterized. Among the few annotated shared genes, *ETHYLENE-RESPONSIVE ELEMENT BINDING FACTOR 13* (*ERF13*) and *RING1*, which have been previously reported to be stress-responsive (Chen *et al*., 2024; Lee *et al*., 2011) were upregulated in HMF (Supplementary Fig. 9A-D).

Taken together, our transcriptomic data suggest that HMF exposure leads to moderate transcriptomic reprogramming, largely in stress-responsive pathways.

### HMF reprograms pathways from growth to defense during seed germination

The transcriptomic results above indicate that HMF is a weaker factor influencing gene expression, compared to germination timing. This promoted us to analyze whether plants respond to the HMF environment in a consistent or varied manner with regard to biological processes. To this end, we first analyzed expression patterns of all DEGs across the three time points using the Short Time-series Expression Miner (STEM) software. In both GMF and HMF conditions, DEGs were predominantly clustered into profile 5 (stable expression followed by an increase) and profile 8 (continuous upregulation over time) (Fig. 4A). A comparative analysis revealed that profile 5 contained 62 genes common to both conditions, with 73 unique to the control and 79 unique to HMF. In profile 8, five genes were shared, while 42 and 45 were specific to CK and HMF, respectively (Fig. 4B). This condition-specific enrichment indicates that HMF has a specific effect on the genes unique to HMF within profile 5 and 8. Gene Ontology (GO) enrichment analysis showed that HMF-specific genes in profile 5 were associated with responses to reactive oxygen species (ROS) and defense against fungi, whereas those in profile 8 were linked to hypoxia response (Fig. 4C, D). In contrast, GMF-specific genes were enriched for different functions, such as cell wall organization and root hair elongation (Fig. 4C). Taken together, these data indicate that HMF induces the dynamic upregulation of unique gene sets associated with hypoxia and ROS responses during germination.

**Fig. 4.**
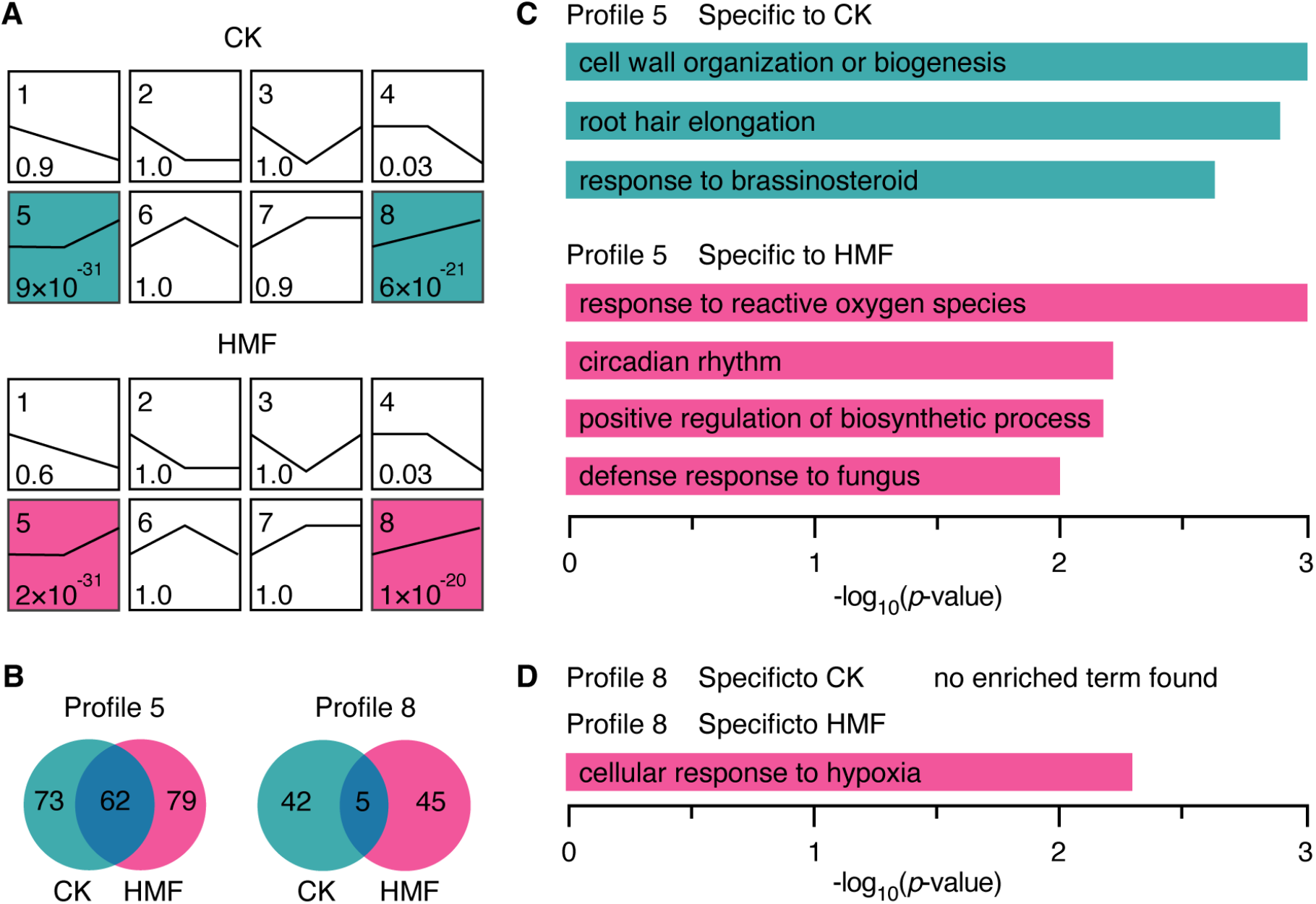
Temporal dynamic expression patterns of DEGs. (A) Short Time-series Expression Miner (STEM) analysis of DEGs under CK and HMF conditions. Each box represents a distinct expression profile, with the number of profiles and *p*-values shown. Profiles 5 and 8 are significantly enriched for both CK and HMF group. (B) Venn diagrams showing the overlap of DEGs in profiles 5 and 8 between CK and HMF groups. (C, D) Gene Ontology (GO) enrichment analysis of CK-and HMF-specific genes in profiles 5 (C) and 8 (D).

We then performed GO enrichment analysis on the HMF-regulated DEGs across the three time points. The transcriptional landscape showed a rapid and sustained activation of defense-related pathways under HMF exposure (Fig 5A-C). At the early stage (28 h), a broad-spectrum stress response was initiated, characterized by significant enrichment of genes responsive to both biotic (bacterium and fungal origin molecule) and abiotic (hyperosmotic salinity, oxidative stress, and cold) stimuli. Concurrently, signaling pathways associated with key stress-related phytohormones, including abscisic acid (ABA), ethylene, and brassinosteroids (BRs), were activated. This defense program was intensified at 36 h and 44 h, broadening to encompass a wider range of environmental challenges. Notably, biosynthetic pathways for sphingolipids, flavonols, and anthocyanins, which have been well-recognized to play an important role in stress adaptation and redox homeostasis, were upregulated at 28 h or 44 h. Furthermore, the emergence of GO terms related to developmental arrest, such as maintenance of seed dormancy and negative regulation of growth at 36 h or 44 h, suggests a compensatory growth limitation in response to HMF-induced stress.

**Fig. 5.**
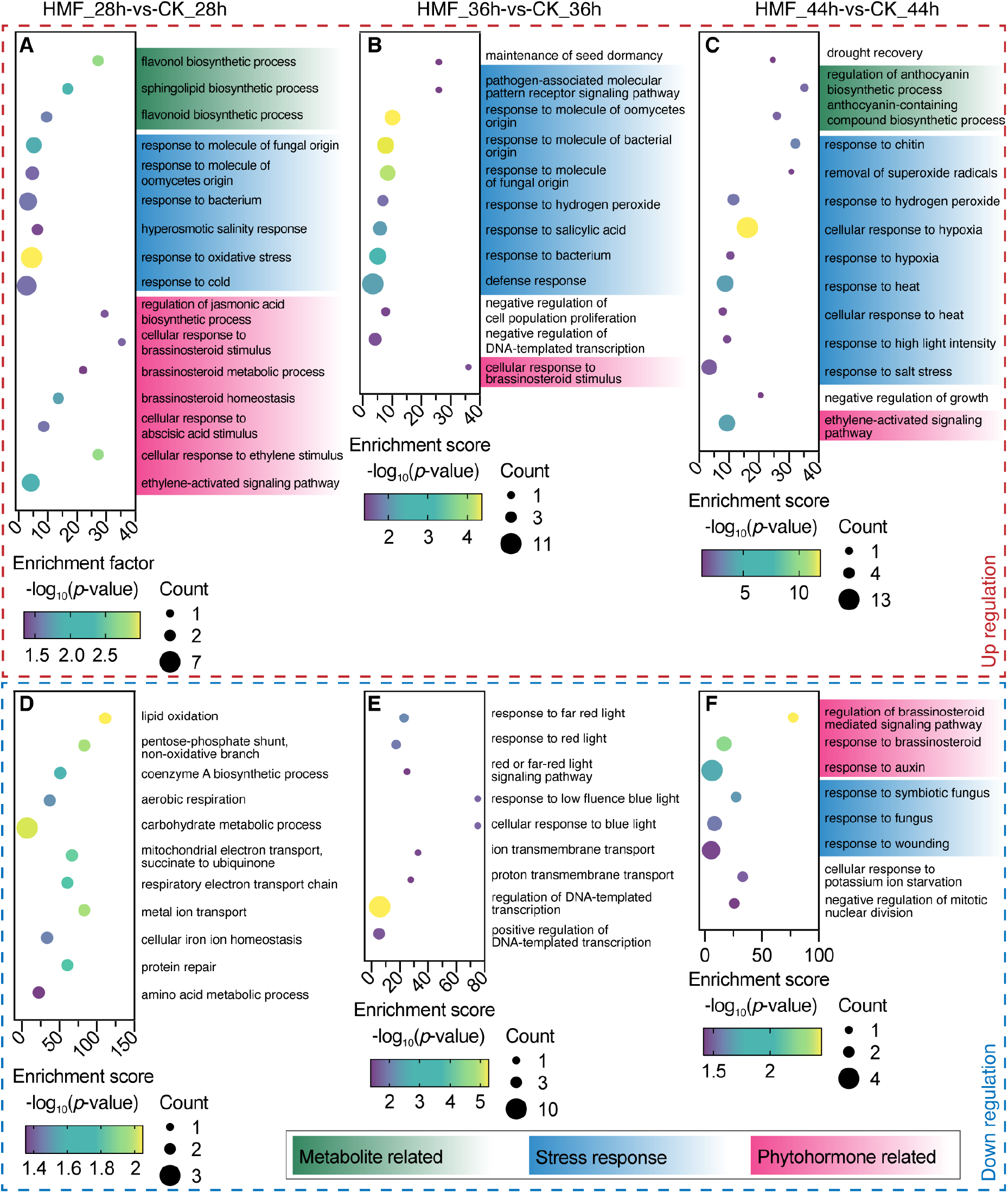
GO enrichment analysis of DEGs at different times of HMF treatment. (A–C) GO enrichment analysis of upregulated DEGs at 28 h (A), 36 h (B), and 44 h (C). Each bubble represents a significantly enriched biological process. Bubble size indicates the number of DEGs in each GO term (Count), color represents statistical significance (-log_10_(*p*-value)), and the x-axis shows the enrichment score. Enriched terms are grouped into functional categories: metabolism-related (green), stress response (blue), and phytohormone related processes (pink). (D–F) GO enrichment analysis of downregulated DEGs by HMF.

Concomitantly, a shutdown of several important metabolic and developmental pathways was evident from the analysis of down-regulated genes (Fig 5D-F). This resource-conserving strategy was initiated early (28 h) with the suppression of central energy metabolism, including aerobic respiration, the mitochondrial electron transport chain, lipid oxidation and carbohydrate metabolism. The biosynthetic pathways for amino acids and coenzyme A were also significantly repressed, further limiting the supply of energy and biosynthetic precursors required for germination. By the mid-stage (36 h), this suppression extended to signaling pathways that promote growth, most notably through the down-regulation of responses to light signals (response to red and blue light). This suggests an active coordinated process to adapt HMF cues. At 44 h, the down-regulation of auxin response indicated a sustained suppression of growth-related processes. Interestingly, the concurrent attenuation of certain stress-responsive pathways may reflect feedback regulation following prolonged stress response, potentially serving to prevent excessive energy expenditure and restore cellular homeostasis.

Finally, we performed a gene set enrichment analysis (GSEA) to identify coordinated changes in predefined biological pathways without relying on significant changes in individual genes. Across all three time points, gene sets related to biotic and abiotic stress responses, including responses to molecules of fungal, oomycete, and bacterial origin molecules, as well as hydrogen peroxide and hypoxia, were consistently enriched under HMF treatment (Supplementary Fig. 10A-H, J, K). These results suggest an early and sustained activation of generalized stress signaling pathways. In contrast, the starch catabolic process was negatively enriched (Supplementary Fig. 10A, I), indicating a possible suppression of energy mobilization during germination under HMF conditions.

Collectively, our deepmining of transcriptomic data reveals that HMF-delayed seed germination is associated with a set of upregulated genes involved in oxidative stress as well as of downregulated genes in growth.

### Moderate ROS accumulation mediates HMF-induced germination delay and primes early stress tolerance

To test the hypothesis that ROS serves as a mediator for HMF-induced germination delay, we assessed ROS accumulation in HMF-and GMF-grown seedlings using NBT (superoxide anion) and DAB (hydrogen peroxide) staining. Our results showed that roots of seedlings exposed to HMF, particularly their tips, were more deeply stained by both NBT and DAB than to GMF (Fig. 6A, C). Staining quantification analysis showed that depth of staining extracted from HMF-treated seedlings were 9.56% (NBT staining) and 8.69% (DAB staining) higher than that from control (Fig. 6B, D). These results suggest that HMF induces significant ROS accumulation during early seedling development.

**Fig. 6.**
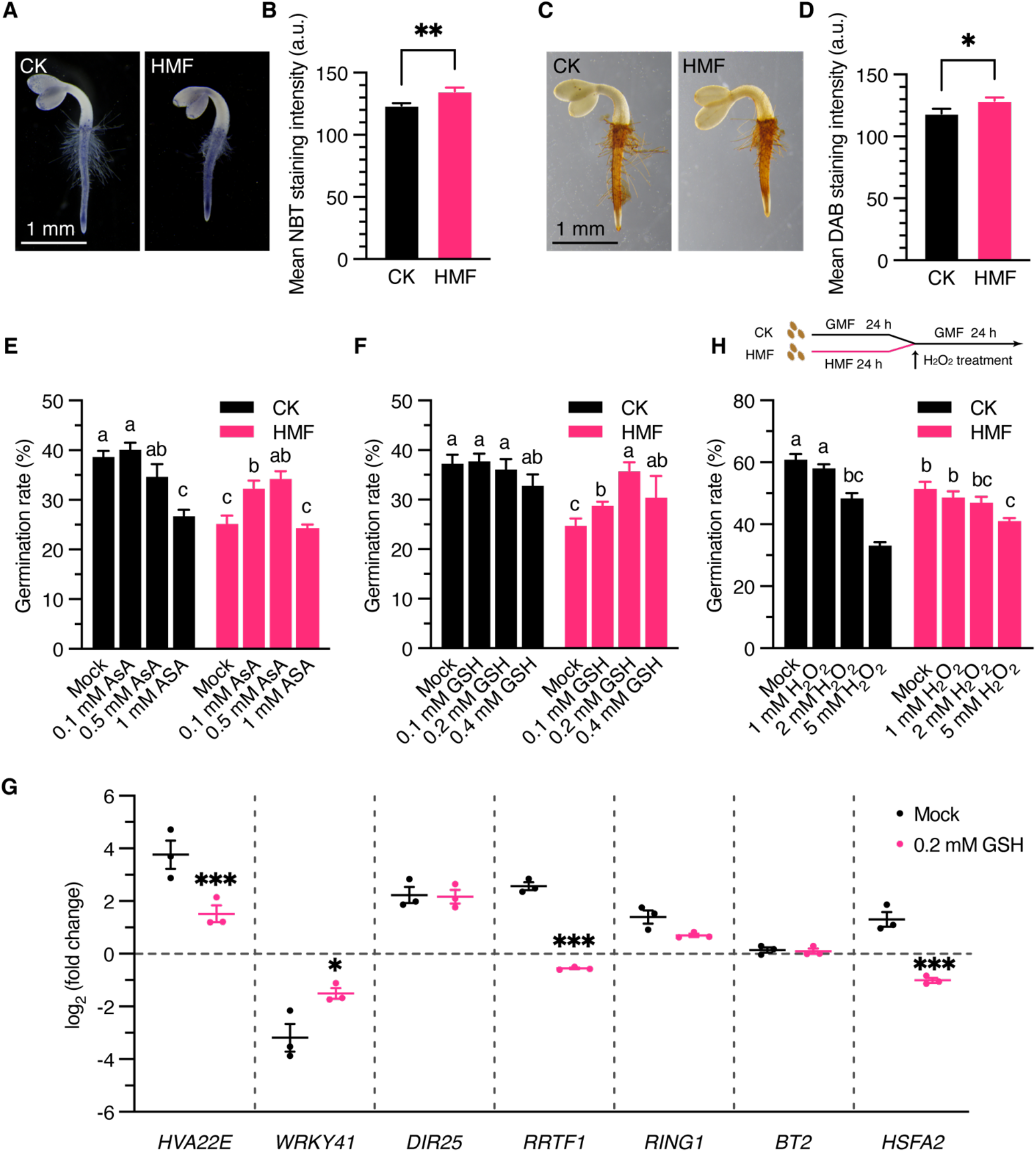
Analysis of ROS roles in HMF-delayed seed germination. (A, B) NBT staining (A) and quantification (B) of ROS in seedlings. Data are shown means ± SE (n ≥ 10). Stars indicate significant differences between CK and HMF (Student’s *t*-test). **, *p* < 0.01. Scale bar, 1 mm. (C, D) DAB staining (A) and quantification (B) of ROS in seedligns. Data are shown means ± SE (n ≥ 11). Stars indicate significant differences between CK and HMF (Student’s *t*-test). *, *p* < 0.05. Scale bar, 1 mm. (E, F) Effects of exogenous AsA (E) and GSH (F) on seed germination at 44 h in HMF and GMF conditions. Data are shown means ± SE (n = 4). Groups that do not share any letter are significantly different (one-way ANOVA followed by Tukey’s post hoc test, *p* < 0.05). (G) Expression of stress-responsive genes in seedlings treated with or without exogenous GSH. Data are shown means ± SE (n = 3). Stars indicate significant differences between CK and HMF (Student’s *t*-test). *, *p* < 0.05; ***, *p* < 0.001. (H) Effects of exogenous hydrogen peroxide on HMF pretreated seeds. Following a 24 h pretreatment under GMF or HMF conditions, seeds were exposed to hydrogen peroxide, and germination rates were evaluated 24 h later. Data are shown means ± SE (n = 4). Groups that do not share any letter are significantly different (one-way ANOVA followed by Tukey’s post hoc test, *p* < 0.05).

To determine whether oxidative stress contributes to the germination delay under HMF conditions, we performed antioxidant rescue experiments with ascorbic acid (AsA) and glutathione (GSH). In GMF, 0.1 mM AsA treatment slightly improved germination at 44 h, whereas higher concentrations (0.5 and 1 mM) delayed germination (Fig. 6E). In contrast, in HMF, 0.1 mM AsA significantly increased germination rate from 25.1% (mock) to 32.0%, and 0.5 mM AsA further increased it to 34.4%, while 1 mM AsA had no effect. Similarly, GSH treatments had minimal or slightly negative effects in GMF (Fig. 6F), whereas in HMF, 0.1 mM GSH slightly improved germination, with greater rescue at 0.2 and 0.4 mM (Fig. 6F). These data suggest that appropriate antioxidant levels can fully counteract HMF-induced germination delays, supporting a key role for ROS in this process.

To investigate whether ROS scavengers alleviate germination delay by modulating stress-responsive genes in HMF, we examined expression of several ROS-related genes selected from DEGs. The qRT-PCR analysis showed that exogenous GSH markedly suppressed the HMF-induced expression of *HVA22E*, *WRKY41*, *RRTF1*, and *HEAT SHOCK TRANSCRIPTION FACTOR A2* (*HSFA2*), while had a no effect on expression of *DIR25*, *RING1*, and *BTB AND TAZ DOMAIN PROTEIN 2* (*BT2*) (Fig. 6G). These results provide molecular basis supporting that GSH which significantly rescued germination in HMF is involved in seedling defense induced by ROS. Intriguingly, seeds pretreated with HMF for 24 hours exhibited reduced sensitivity to exogenous H_2_O_2_. In GMF, low concentrations of exogenous H_2_O_2_ had minimal impact on seed germination, whereas higher concentrations progressively inhibited germination (Fig. 6H). In contrast, HMF pretreated seeds exhibited more tolerance to H_2_O_2_ than the GMF control (Fig. 6H). Together, these findings suggest that HMF-induced stress responses prime seeds for enhanced tolerance to subsequent oxidative stress during germination.

### HMF delays seed germination through both CRY-dependent and -independent pathways

To investigate whether cryptochrome (CRY) signaling, a proposed mediator for magnetoreception (Xie, 2022), is involved in the HMF-induced delay of seed germination, we compared the germination rate of wild-type (WT) and *cry1 cry2* double mutant seeds under both white and blue light conditions. Under white light, both WT and *cry1 cry2* seeds exhibited a comparable delay in germination in HMF, with significantly reduced germination rates at 44 and 46 h after sowing (Fig. 7A, B), indicating that the delay in white light is largely independent of CRY1/CRY2. In contrast, under blue light, WT seeds germinated similarly under HMF and GMF at all examined times with exception of at 44 h (Fig. 7C), whereas *cry1 cry2* mutants were completely insensitive to HMF (Fig. 7D). These findings indicate that the HMF effect on seed germination depends on light quality: CRYs mediate the response under blue light, while a CRY-independent pathway operates under white light.

**Fig. 7.**
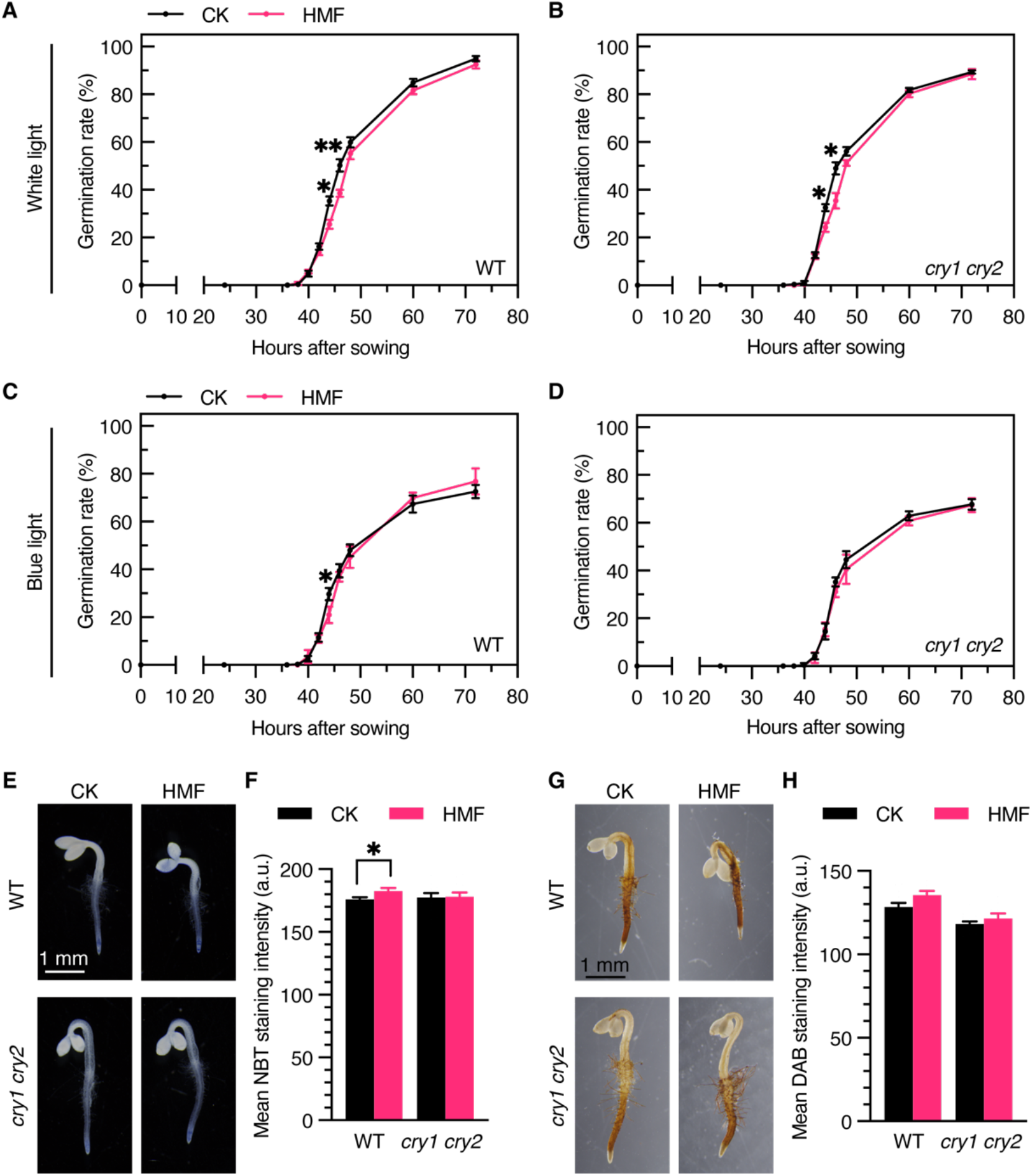
HMF-induced germination delay is partially dependent on CRY signaling. (A, B) Germination assay of WT (A) and *cry1 cry2* (B) seeds in white light. Data are shown means ± SE (n = 4). Stars indicate significant differences between CK and HMF (Student’s *t*-test). *, *p* < 0.05; **, *p* < 0.01. (C, D) Germination assay of WT (C) and *cry1 cry2* (D) seeds in blue light. Data are shown means ± SE (n = 4). *, significant difference between CK and HMF (Student’s *t*-test, *p* < 0.05). (E, F) *In situ* NBT staining (E) and quantification (F) of ROS in WT and *cry1 cyr2* seedlings grown in blue light. Data are shown means ± SE (n ≥ 14). Stars indicate significant differences between CK and HMF (Student’s *t*-test). *, *p* < 0.05. Scale bar, 1 mm. (G, H) *In situ* DAB staining (G) and quantification (H) of ROS in WT and *cry1 cyr2* seedligns grown in blue light. Data are shown means ± SE (n ≥ 10).

To explore the relationship between CRYs and HMF-induced ROS changes, we examined the ROS levels in WT and *cry1 cry2* seedlings exposed to HMF and GMF. Under white light, both genotypes showed significantly stronger ROS staining (NBT and DAB) in HMF than in CK (Supplementary Fig. 11). However, under blue light, NBT- and DAB-staining intensities were generally lower in both HMF- and GMF-treated seedlings, particularly in their tips (Fig. 7E, G), compared to under white light, suggesting that besides blue light other light spectra are also involved in HMF-induced ROS accumulation. Quantification analysis of ROS content detected by NBT staining was significantly higher in HMF than in GMF, whereas that by DAB staining did not differ significantly (Fig. 7F, H). In contrast, both NBT- and DAB-staining in *cry1 cry2* was comparable between CK and HMF (Fig. 7E-H). These results suggest a role of CRYs in modulating HMF-induced ROS accumulation under blue light.

To explore potential CRY-independent regulatory mechanisms, we constructed gene regulatory networks (GRNs) from transcriptomic data using GENIE3, a machine learning-based algorithm for inferring regulatory interactions. To enhance biological interpretability, we restricted the regulator candidates to DEGs and annotated transcription factors from PlantTFDB, rather than the full set of expressed genes. By retaining only regulatory edges involving DEGs, we reconstructed sub-GRNs specific to the HMF at three distinct time points. However, the resulting networks did not exhibit clear hierarchical structure or dominant regulatory hubs (Supplementary Fig. 12-14). Instead, the networks appeared highly fragmented and decentralized, with the majority of regulators regulating only one or two target genes. This dispersed topology may be attributed to the relatively small number of DEGs identified under HMF, which limits the resolution of network inference.

To identify potential core regulators consistently active across germination process, we next focused on regulators that were predicted to exert regulatory influence at all three time points. This analysis yielded 14 shared regulatory genes, including eight transcription factors (Supplementary Fig. 15A). Among these, several were previously associated with stress responses including *CALMODULIN LIKE 37* (*CML37*), *SALT TOLERANT CALLUS 8* (*STC8*), *HSP17.6A*, and *SPEECHLESS* (*SPCH*) (Ahmad *et al*., 2015; Changle *et al*., 2006; Scholz *et al*., 2015; D. Wang *et al*., 2025), while others were involved in seed germination processes including *EARLY-PHYTOCHROME-RESPONSIVE1* (*EPR1*) and *PHYTOCHROME-INTERACTING FACTOR* 6 (*PIF6*) (Liu *et al*., 2021; Penfield *et al*., 2009). PIF6, a known light-responsive transcription factor, drew our attention due to its established role in seed germination and its regulation through alternative splicing. Previous studies have shown that the *PIF6-α* isoform has limited impact on germination, whereas the *PIF6-β* isoform promotes germination (Penfield *et al*., 2009). Although the overall transcription level of *PIF6* was not significantly altered under HMF, transcript isoform expression analysis revealed isoform-specific dynamics: the α transcript showed a non-significant reduction at 44 h, while the β transcript was significantly downregulated at 28 h but markedly upregulated at 44 h (Supplementary Fig. 15B, C). These findings suggest that under HMF, PIF6 may undergo a functional switch from repress to promotion in seed germination, potentially contributing to the observed delay rather than complete inhibition of germination process.

## Discussion

In this study, we investigated HMF effect on seed germination and the underlying molecular mechanisms. Our results demonstrate that HMF delays seed germination by inducing a moderate but significant accumulation of ROS. This delay is partially counteracted by supplementing the growth media with antioxidants such as GSH and AsA, confirming a causal link between ROS accumulation and the germination phenotype. Interestingly, seeds imbibed under HMF acquired a primed tolerance to subsequent oxidative stress, suggesting that HMF-induced ROS function as signaling molecules to enhance stress preparedness. These physiological observations are strongly supported by transcriptomic profiling, which revealed that HMF exposure reprograms gene expression, shifting the balance from growth-promoting to defense-related pathways during early germination (Fig. 8).

**Fig. 8.**
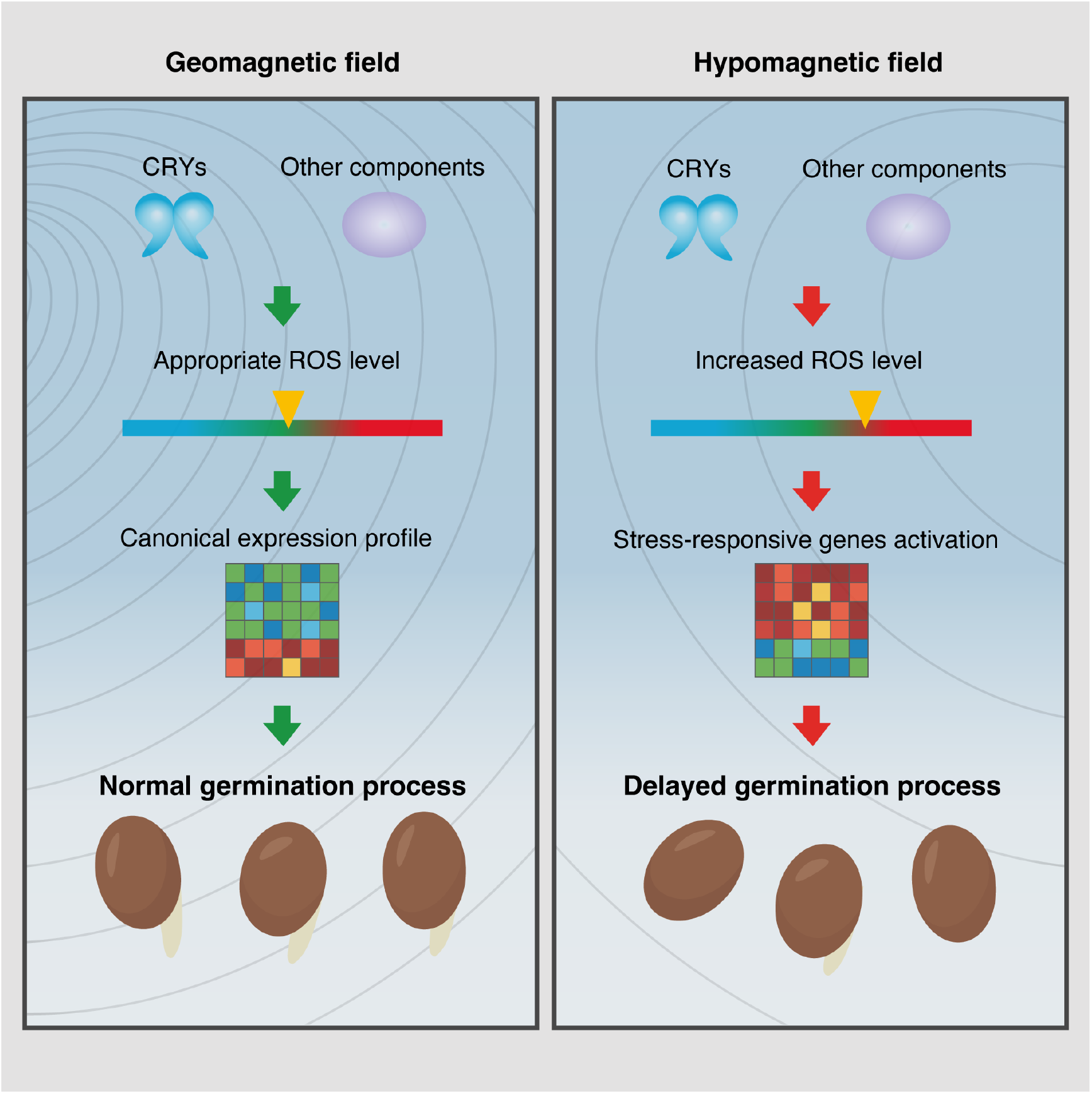
The possible mechanism underlying HMF-induced germination delay. Under geomagnetic field, appropriate levels of ROS promote seed germination. In contrast, exposure to hypomagnetic field induces moderate ROS accumulation in a CRY-dependent and -independent manner, which activates the expression of stress-responsive genes and drives transcriptional reprogramming. This altered gene expression profile ultimately results in delayed germination process.

Beyond delaying seed germination, HMF exposure also inhibited primary root growth and cotyledon expansion, in line with previous reports (Mo *et al*., 2011; Xu *et al*., 2012). Although ROS are well documented to play dual roles in both seed germination and post-germinative growth and development (P. Wang *et al*., 2024; Y. Wang *et al*., 2025), it remains unclear whether the growth retardation observed under HMF conditions is also mediated by ROS. Parmagnani *et al*. (2022) reported lower H_2_O_2_ levels in seedlings grown in HMF, suggesting that the effect of HMF on ROS homeostasis may differ on developmental stages. This stage-dependent effect is consistent with the multiple roles of ROS in plants (Gogoi *et al*., 2024; Mittler *et al*., 2022). Under abiotic or biotic stress conditions, moderate ROS accumulation can trigger defense gene expression, activate antioxidant systems, and modulate hormone signaling pathways such as those of ABA and salicylic acid (Gogoi *et al*., 2024; Mittler *et al*., 2022), whereas excessive ROS can lead to oxidative damage and programmed cell death (Castro *et al*., 2021; Mittler, 2017). In germinating seeds, ROS gradients are essential for cell proliferation, cell wall loosening, and endosperm weakening (Müller *et al*., 2009; Oracz and Karpinski, 2016), and any perturbation of this delicate balance, as imposed by HMF, could delay the transition from quiescence to active growth. In this study, HMF induced a moderate increase in ROS levels that delayed the process of seed germination, but did not affect the final germination rate. These findings suggest that HMF-induced ROS accumulation remains within the physiological signaling range, serving to initiate adaptive stress responses. This interpretation is consistent with our results that exogenous GSH treatment, which quenches ROS, reverses the phenotype of delayed seed germination and normalizes the expression of a subset of stress-responsive genes in HMF (Fig. 6G). Similarly, the primed oxidative stress tolerance observed in HMF-imbibed seeds is reminiscent of the well-characterized priming phenomenon, whereby initial exposure to a mild stress enhances tolerance to subsequent, more severe stress events (Hossain *et al*., 2015; Savvides *et al*., 2016).

ROS homeostasis is regulated by the balance between production and scavenging pathways. Our transcriptomic data revealed a significant upregulation of genes associated with ROS generation. For example, *RESPIRATORY BURST OXIDASE HOMOLOG A* (*RBOHA*), encoding a plasma-localized NADPH oxidases, was significantly upregulated at 36 h under HMF. NADPH oxidases are central mediators of the apoplastic ROS burst and have been shown to play critical roles in systemic signaling, cell-to-cell ROS wave propagation, and stress acclimation (Suzuki *et al*., 2011; Zandalinas *et al*., 2020; Mhamdi, 2022). The upregulation of *RBOHA* under HMF therefore provides a mechanistic basis for the observed ROS accumulation. In contrast, the transcript levels of key ROS-scavenging enzymes, including superoxide dismutases, catalases, and peroxidases, remained unchanged (Supplementary Fig. 16), indicating that the enhanced ROS levels under HMF are primarily driven by increased production rather than impaired scavenging. In addition, activities of ROS metabolic enzymes are also post-translationally regulated by magnetic fields (Tao *et al*., 2025; S. Wang *et al*., 2024). Taken together, it is likely that HMF regulates ROS levels through multiple mechanisms. It is noteworthy that many of the top DEGs induced by HMF remain functionally unclear. Characterizing these unknown genes in the future may reveal additional components of the HMF-triggered ROS signaling cascade in plants.

The radical pair mechanism (RPM), in which blue-light photoreceptors cryptochromes form spin-correlated radical pairs whose singlet-triplet interconversion kinetics can be modulated by external magnetic fields, represents the most well-characterized model for biological magnetoreception (Hore and Mouritsen, 2016). This mechanism has been extensively studied in animal (Xie, 2022), and also implicated in plant magnetosensing (Thoradit *et al*., 2023). *Arabidopsis* mutants lacking CRYs display insensitivity to magnetic field variations in a range of physiological processes including root growth, hypocotyl elongation, anthocyanin accumulation and flowering time (Agliassa *et al*., 2018; Ahmad *et al*., 2007; Jin *et al*., 2019; Xu *et al*., 2012). However, it remains unclear whether HMF-delayed seed germination is mediated through a CRY-dependent pathway. Our results showed that the inhibitory effect of HMF on seed germination is completely abolished in *cry1 cry2* under blue light, confirming a CRY-dependent pathway. However, the germination delay was still remained under white light even in CRY-deficient mutants (Fig. 7), suggesting the presence of a CRY-independent pathway. Consistently, the HMF-induced ROS change followed the same pattern as the germination phenotype (Fig. 7E-H), reinforcing the tight coupling between ROS and germination control under HMF. Indeed, our further analysis revealed that the CRY-independent pathway for the HMF-induced ROS accumulation is related to *PIF6* and *EPR1*, known to participate in red light–regulated seed germination (Supplementary Fig. 8). This observation raises a possibility that phytochrome signaling may integrate with magnetosensing in a manner independent of classical RPM. Future studies should focus on identifying additional potential magneto-sensitive components, such as iron-sulfur proteins, ion channels, and non-CRY photoreceptors, to build a comprehensive understanding of plant magnetoreception.

Over geological timescales, GMF has undergone multiple phases of weakening and polarity reversal with excursions and transitions lasting thousands of years (Valet and Fournier, 2016). During periods of low geomagnetic intensity, the reduced magnetospheric shielding allows more cosmic radiation to reach the Earth, potentially increasing stress in terrestrial organisms. Some studies have speculated on a link between geomagnetic fluctuations and key evolutionary events, such as the diversification and rapid radiation of angiosperms (Occhipinti *et al*., 2014). In this evolutionary context, the ability to detect geomagnetic weakening and mount a preemptive stress response would confer a significant selective advantage. Our finding that HMF induces moderate ROS accumulation that is sufficient to delay germination and prime stress tolerance is consistent with such an adaptive scenario. Thus, the ROS-mediated response to HMF may represent a conserved environmental sensing mechanism that has been shaped by the evolutionary trajectory of GMF.

## Methods

### Plant materials and growth conditions

The wild-type *Arabidopsis thaliana* used in this study was the Columbia-0 (Col-0) ecotype. The *cry1 cry2* used was consistent with previous study (Jin *et al*., 2019). Seeds were surface-sterilized and stratified at 4℃ in darkness for 48 h prior to sowing on half strength Murashige & Skoog (MS) medium supplemented with 1% sucrose. Seedlings were grown at 22℃ under long-day conditions (16 h light/8 h dark) with white light at an intensity of 100 μmol ·m^-2^·s^-1^. For blue light cultivation, seeds were grown under continuous light with intensity of 30 μmol ·m^-2^·s^-1^.

For germination assay, seeds were directly placed in HMF or GMF environment after sowing. For root length and cotyledon area measurements, seeds were germinated and grown under the GMF for 4 days, then transferred to either GMF or HMF conditions for an additional 3 days growth before data collection. For flowering time assays, seeds were grown under the GMF for 5 days, then transplanted into soil and maintained under GMF or HMF conditions.

### Setup of hypomagnetic field environment

To establish a HMF environment, a three-dimensional Helmholtz coil system was employed. The system consists of three orthogonally arranged square coils (side length of 120 cm for x-axis, 124.5 cm for y and 129 cm for z) that generate compensatory magnetic fields to counteract the GMF. A triaxial fluxgate magnetometer (9200B, Shanghai Haibi, range of -99, 999 to 99, 999 nT, ±1 nT resolution, 1 Hz sampling rate) continuously monitors the magnetic field, while a control software (FE-2100FG, Hunan forever elegance Technology Co., Ltd, Loudi, Hunan, China) dynamically adjusts the current input to maintain the HMF condition every 5 s. Generally, the samples were kept in center cubic space with side length of 10 cm (below 0.2 μT when center is near zero in HMF).

The control group was set in an identical coil system without current input, thereby maintaining the ambient GMF. For each experiment, the coil set used to generate the HMF was randomly selected to minimize systematic bias. To ensure environmental consistency, all systems were placed in a separate, light-shielded, temperature-controlled room. The representative field record is showed in Supplementary Fig. 2 and Supplementary Table 3 and 4.

Furthermore, we also employed a passive shielding device to create HMF. The passive shielding device (Hefei Moce Technology Co., Ltd., Hefei, Anhui, China) is consisted of four high-permeability material (permalloy 1J85) layers. The inside magnetic field was lower than 30 nT during installation and the running condition is verified by a portable fluxgate magnetometer (TM4100B, Tunkia Co., Ltd., Changsha, Hunan, China; range of -1000 to 1000 μT, ±0.1 μT resolution) with intensity of 0.0 μT (lower than detection limit).

### Assessment of growth and developmental parameters

Seed germination was defined as the emergence of the radicle through the seed coat. At each time point, germination rates were assessed using four replicates, each containing more than 100 seeds. Leaf area and root length were recorded using a stereo microscope (SZX16, Olympus) and an Epson Perfection V800 scanner and quantified with ImageJ (version 1.54d). Bolting time was defined as the number of days from sowing until the first inflorescence stem reached 5 cm in height, while flowering time was recorded as the day of first flower opening.

### RNA-seq library preparation and sequencing

Seeds were harvested at 28 h, 36 h, and 44 h with 3 independent replicates. Total RNA was extracted using the TRIzol reagent (Invitrogen) as described by Zhang *et al*. (2026). RNA purity and concentration were assessed using the NanoDrop 2000 (Thermo Scientific), and RNA integrity was evaluated with the Agilent 2100 Bioanalyzer (Agilent Technologies). RNA-seq libraries were prepared using the VAHTS Universal V6 RNA-seq Library Prep Kit (Vazyme). The mRNA enriched by oligo(dT) magnetic beads, fragmented, reverse-transcribed using the dUTP method for strand-specific library construction. Then library was sequenced by Illumina Novaseq 6000 platform to obtain 150 bp paired-end reads (over 40 million read pairs per sample).

### RNA-seq analysis

Raw reads in FASTQ format were preprocessed using fastp to remove adaptor sequences and low-quality reads. Then the clean reads were mapped to the reference genome (TAIR10.1) by using HISAT2 (v2.1.0). The read counts for each gene were obtained using HTSeq-count (v0.11.2) and gene expression levels were quantified as FPKM. Principal component analysis and distance matrix analysis were performed in R to evaluate the biological replicability among samples. Differentially expressed genes (DEGs) were identified using DESeq2 (v1.48.2) with unadjusted *p* < 0.05 and absolute fold change > 1.5, as multiple testing correction was not applied due to the small magnitude of expression changes. Gene Ontology (GO) enrichment analyses of DEGs were performed in R based on the hypergeometric distribution. Besides, gene set enrichment analysis (GSEA) was performed using clusterProfiler package (v3.10.1) (Subramanian *et al*., 2005; Yu *et al*., 2012).

### Short time-series expression analysis

The median FPKM values of DEGs were used as the input matrix for time-series expression analysis using Short Time-series Expression Miner (STEM) software (Ernst and Bar-Joseph, 2006). The data were log-transformed and clustered using the STEM algorithm. Significant enriched expression profiles (*p* < 0.01) were identified. DEGs assigned to distinct temporal expression clusters were selected for subsequent analyses. Go analyses of DEGs unique to CK and HMF within the same expression profile was performed using Metascape (Y. Zhou *et al*., 2019).

### Quantitative real-time PCR validation

Total RNA was extracted using the RNA Easy Fast Plant Tissue Kit (DP452, TIANGEN). In general, the cDNA was synthesized via PrimeScript RT reagent Kit with gDNA Eraser (Takara). Quantitative real-time PCR validation was performed on QuantStudio 3 (Thermol Scientific) with Hieff qPCR SYBR Green Master Mix (Yeasen). Melting curve analysis was performed following the completion of PCR amplification to verify the specificity of the product. Relative mRNA expression levels were calculated using the 2^−ΔΔCT^ method, with *Actin2* serving as internal reference gene for normalization. For each biological replicate, gene expression was assessed using three technical replicates. The primers were listed in Supplementary Table 5.

### Reactive oxygen species staining and quantification

To detect ROS in *Arabidopsis* seedlings, 3,3’-diaminobenzidine (DAB) and nitroblue tetrazolium (NBT) staining were performed to visualize hydrogen peroxide (H_2_O_2_) and superoxide (O_2_·^−^), respectively. Seeds germinated 60 h after sowing were collected in coils and incubated in either DAB solution (0.5 mg/mL, pH 3.8) or NBT solution (0.1% w/v NBT in 10 mM PBS, pH 6.8) *in situ* or outside of the coils as noted. Staining was performed at room temperature with gentle shaking in the dark for 15 min. Following staining, seedlings were washed and incubated in 95% ethanol overnight for decolorization. Images were acquired using a stereo microscope (SZX16, Olympus) under identical settings. For semi-quantitative analysis, images were converted to 8-bit grayscale and inverted in ImageJ. The mean gray values of seedlings were measured and the raw quantification data were provided in Supplementary Table 6.

### ROS scavengers and H_2_O_2_ treatment

Surface-sterilized seeds were sown on half-strength MS medium supplemented with the indicated concentrations of ascorbic acid (AsA) or glutathione (GSH). Seeds were grown under either the GMF or HMF conditions for 44 h, after which germination rates were determined. For H_2_O_2_ treatment, seeds were first pre-incubated on wet filter paper under GMF or HMF conditions for 24 h, then transferred to GMF conditions with the addition of H_2_O_2_ for another 24 h. Germination rates were subsequently measured.

### Gene regulatory network inference and visualization

The GENIE3 (v1.30.0) (Huynh-Thu *et al*., 2010) was used to construct a transcriptional regulatory network using the complete transcriptome datasets in R. Gene expression levels were quantified as FPKM and log_2_-transformed prior to downstream analyses. Transcription factors which were acquired from PlantTFDB (Jin *et al*., 2017) and DEGs were selected as candidate regulators. All expressed genes were considered as potential targets. A global regulatory network was inferred from the combined expression matrix and regulatory edges with weights >0.01 were retained. Time point-specific subnetworks were subsequently derived by extracting edges whose target genes were DEGs at the corresponding time point.

For visualization, the top 150 edges (ranked by weight) from each subnetwork were displayed using igraph (v2.2.1), tidygraph (v1.3.1) and ggraph (v2.2.2) packages. To illustrate the regulatory dynamics of shared regulators across time points, all associated edges were visualized.

### Transcript isoform expression analysis

RNA-seq clean reads were pseudo-aligned to the *Arabidopsis thaliana* reference transcriptome (Araport11) using Kallisto (v0.51.1) with default parameters (Bray *et al*., 2016). Transcript abundance was quantified in transcripts per million (TPM) and estimated counts, with 100 bootstrap replicates to assess quantification uncertainty. Transcript counts matrix were imported into R and analyzed by DESeq2 (v1.48.2).

## Acknowledgements

We thank Y. Tian (Shanghai Normal University) for technical assistance of ROS detection. We thank Z. Huang (Fudan University), W. Ma (Fudan University) and H. Chen (Center for Excellence in Molecular Cell Science) for providing passive magnetic shielding devices and related technical assistance.

The ChatGPT is used for grammar checking and phrasing suggesting of this manuscript. The Python scripts of magnetic field data collection and visualization are generated with the help of Yuanbao (Tencent). No scientific ideas, data interpretation or images was generated by AI.

## Data availability

The raw RNA-seq data of this study have been uploaded to the SRA (https://www.ncbi.nlm.nih.gov/sra) with accession number PRJNA1393870.

