## Supplemental figures for "Hypomagnetic field inhibits seed germination via ROS signaling and reprogramming transcriptome in *Arabidopsis thaliana*"

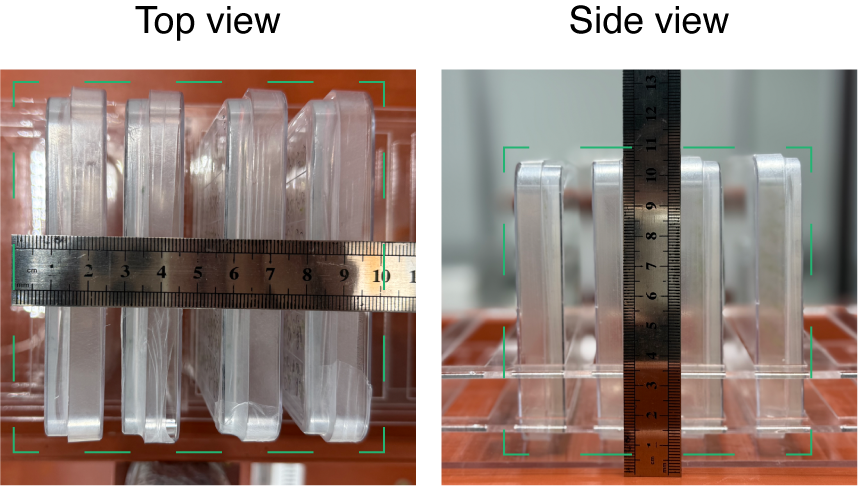

### Supplementary Fig. 1. Typical sample treatment area.

Four technical replicates were confined in a 10 cm side-length region at the center of coils which were marked by green dotted lines.

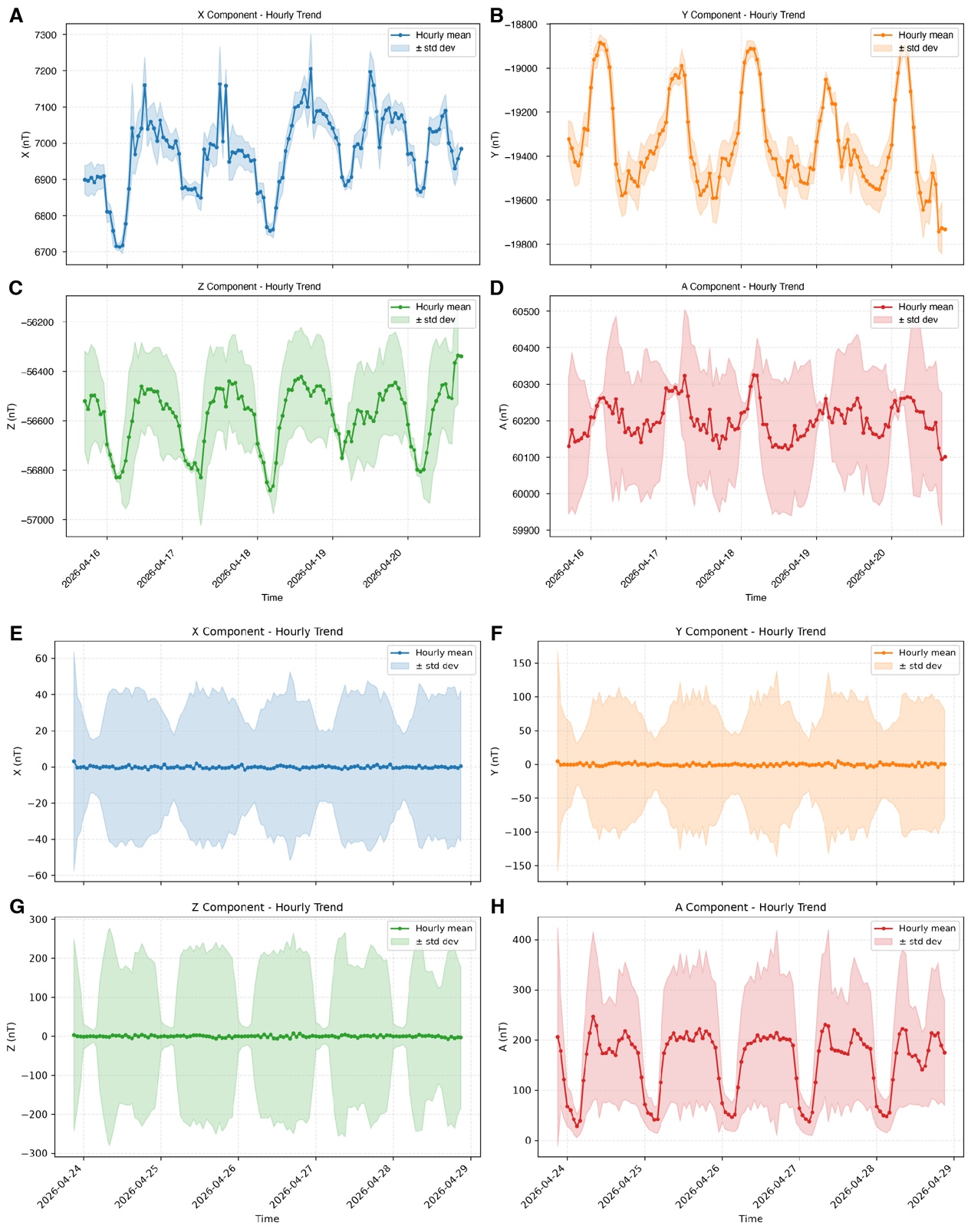

### Supplementary Fig. 2. The representative record of GMF and HMF throughout the experiment.

The field intensity of x, y, z and A (total) are recorded every second throughout the experiment (over 120 h) and visualized by hourly trend. A-D show the GMF data while the E-H show HMF data. The horizontal axis shows the recorded date. Data are shown mean ± SD.

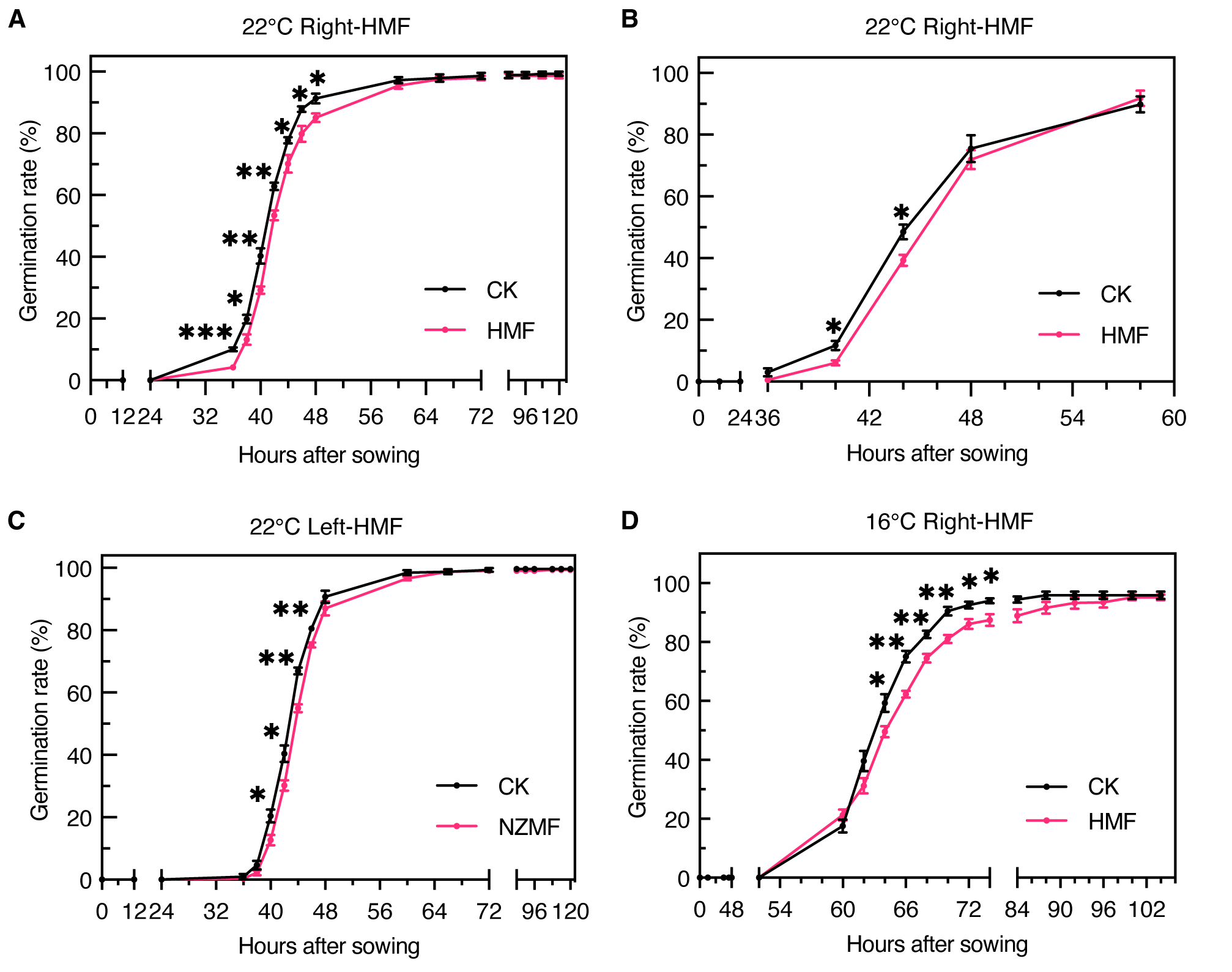

### Supplementary Fig. 3. The repeat data of HMF effects on seed germination.

The experiment temperature and HMF-generating coils are labeled above the picture. Data are shown means ± SE (n = 4). Stars indicate significant differences between CK and HMF (Student’s *t*-test). *, *p* < 0.05; **, *p* < 0.01; ***, *p* < 0.001.

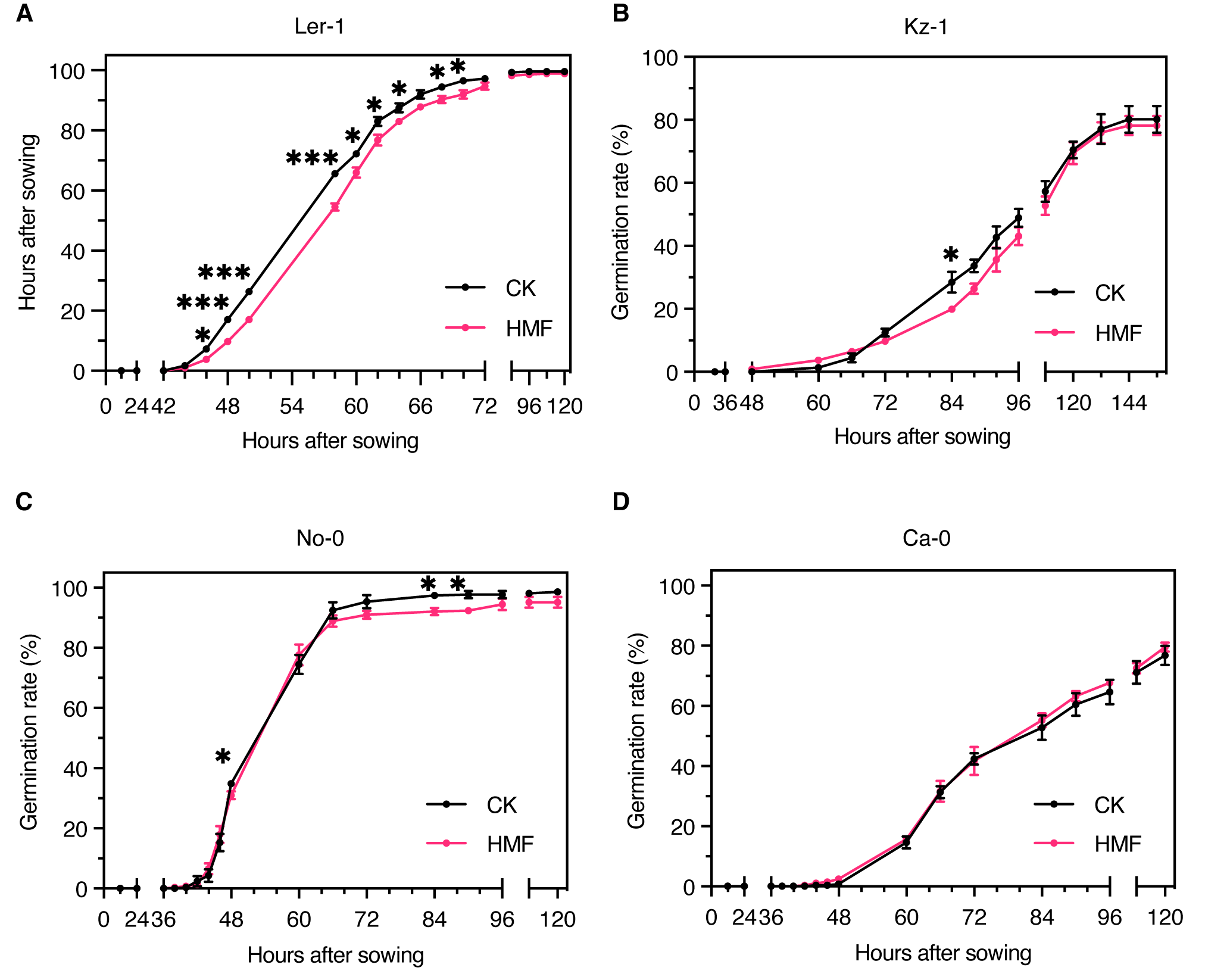

### Supplementary Fig. 4. The effects of HMF on seed germination of different *Arabidopsis* ecotypes.

The name of ecotype is labeled above the picture. Data are shown means ± SE (n = 4). Stars indicate significant differences between CK and HMF (Student’s *t*-test). *, *p* < 0.05; ***, *p* < 0.001.

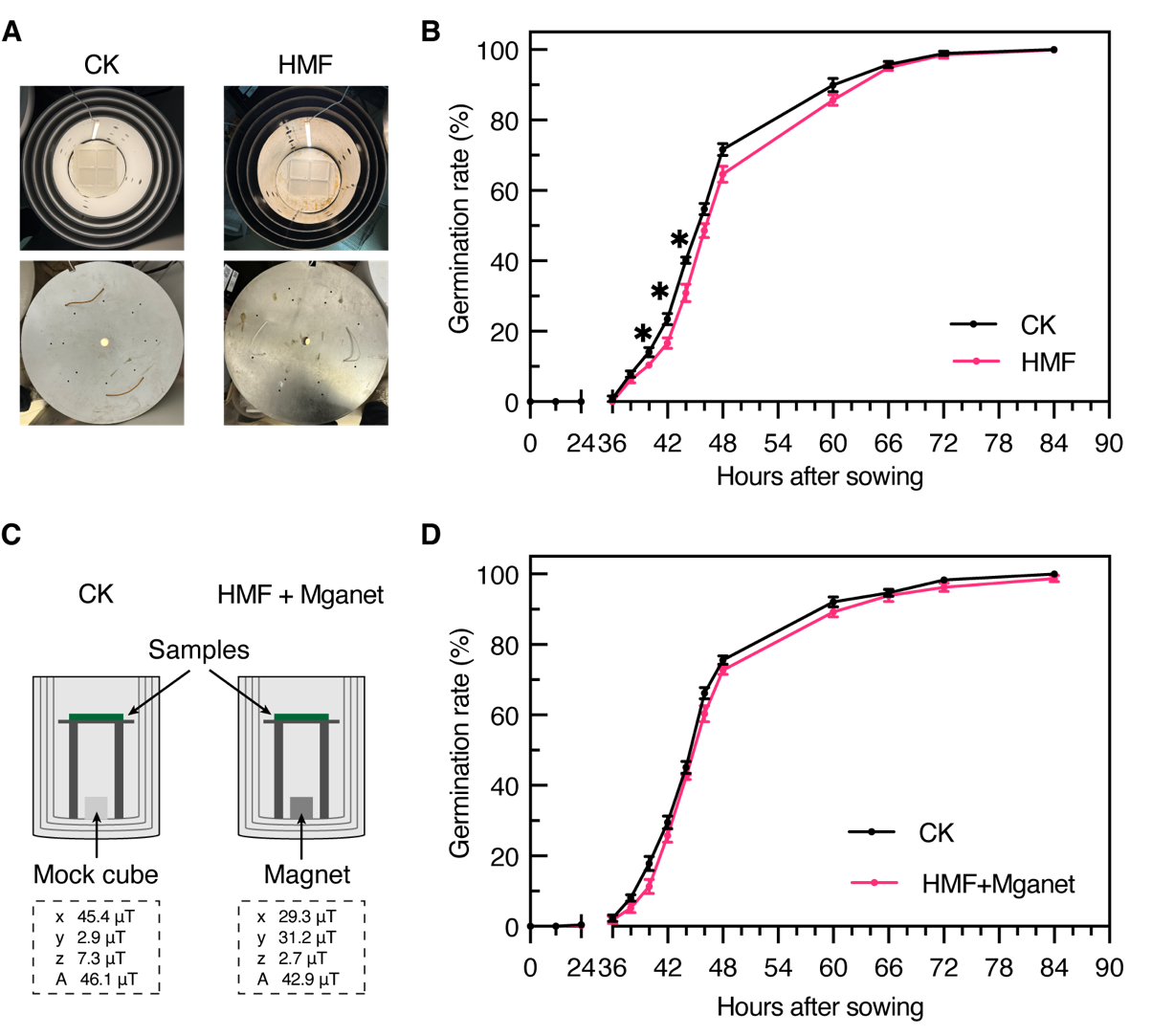

### Supplementary Fig. 5. HMF generated by passive shielding delays seed germination of *Arabidopsis*.

(A) Top views of the passive shielding device (HMF) and its corresponding mock device (CK). The shielding device consists of four layers of high-permeability material, whereas the mock device is made of plastic. Each layer has a lid; the lower image shows the assembled device with lids closed.

(B) HMF delays seed germination. Data are shown means ± SE (n = 4). Stars indicate significant differences between CK and HMF (Student’s *t*-test). *, *p* < 0.05.

(C) Diagram of the magnetic field recovery experiment. The field is recovered by setting a magnet in the shielding device. The filed intensity of sample area is measured and shows below. For CK group, a non-magnetic cube is settled in the mock device.

(D) The magnetic field recovery eliminated the delay in seed germination. Data are shown means ± SE (n = 4). No significant differences between CK and HMF were detected (Student’s *t*-test).

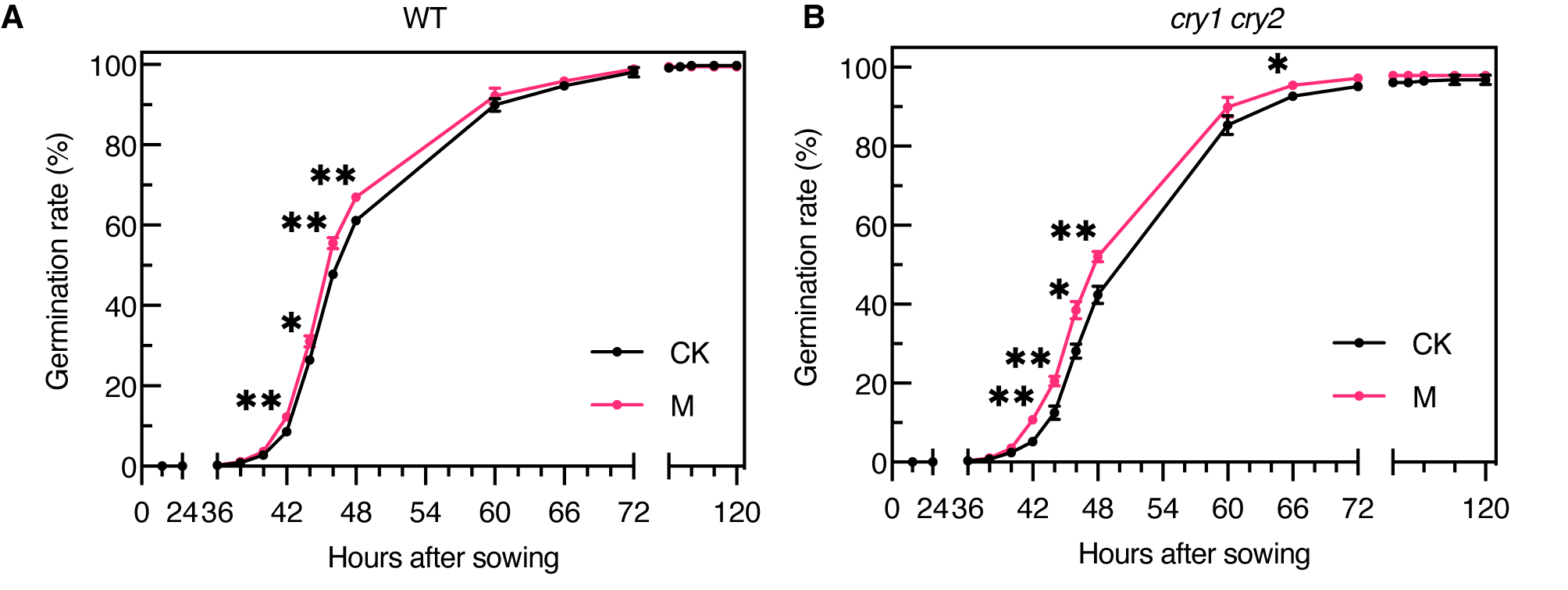

### Supplementary Fig. 6. Static magnetic field of 600 mT promotes seed germination of both WT and *cry1 cry2*.

The static magnetic field is created by a NdFeB magnet. An identical size unmagnetized block is used for CK. Seeds were cultivated under the same condition as in HMF except the field is different. Data are shown means ± SE (n = 4). Stars indicate significant differences between CK and static magnetic field (M) (Student’s *t*-test). *, *p* < 0.05; **, *p* < 0.01.

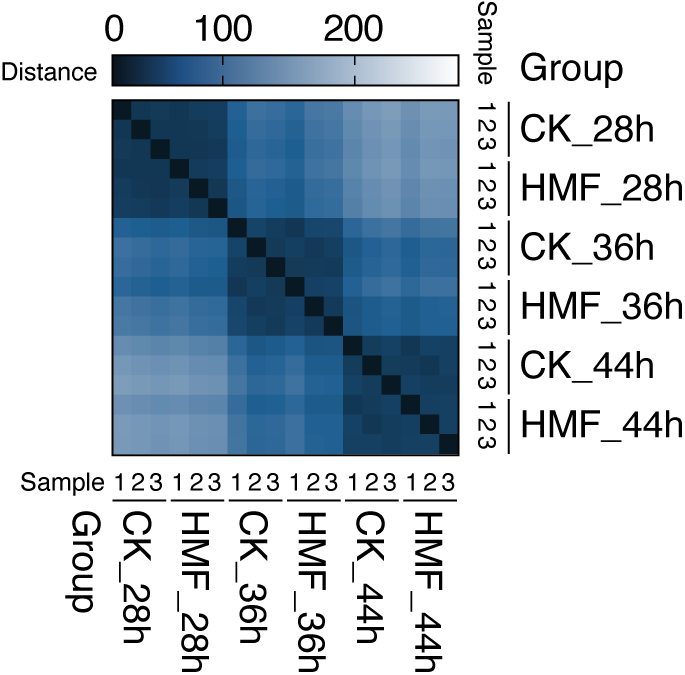

### Supplementary Fig. 7. Heatmap of pairwise sample distances from transcriptomic profiling.

Darker colors (closer to 0) indicate smaller distances and thus higher similarity between samples.

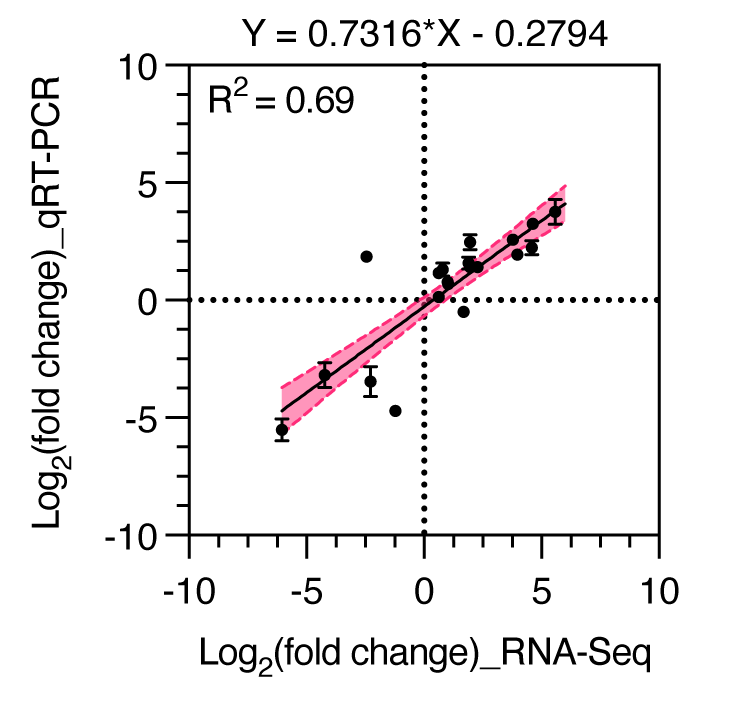

### Supplementary Fig. 8. Validation of DEGs by qRT-PCR.

Correlation analysis between RNA-Seq and qRT-PCR results. The x-axis shows log_2_ (fold change) values from RNA-Seq, and the y-axis shows corresponding values from qRT-PCR (mean ± SE). Each dot represents a validated gene. The pink shaded area represents the 95% confidence interval of fitted curve.

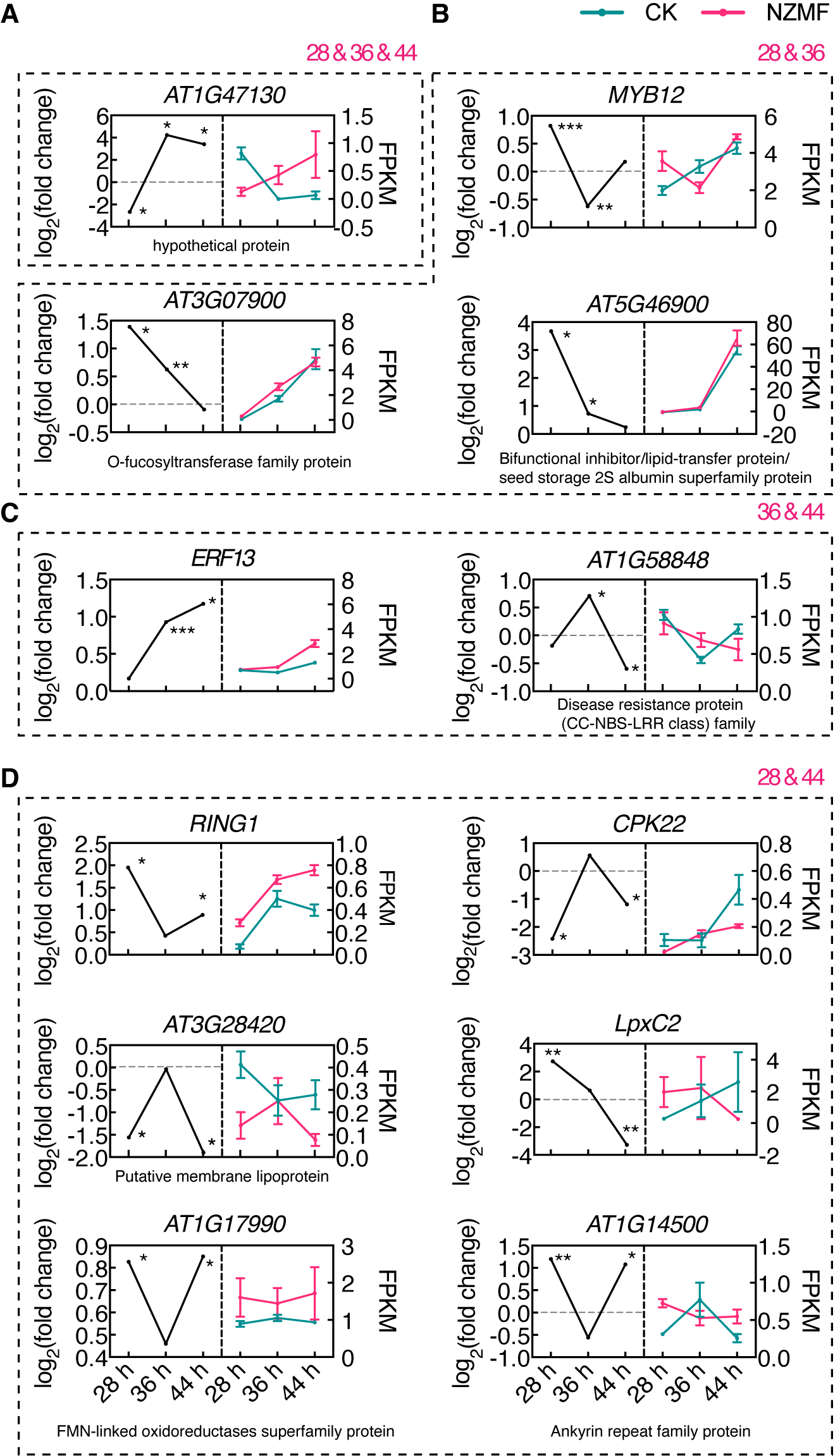

### Supplementary Fig. 9. Time-dependent expression trends of overlap DEGs.

(A-D) Expression pattern of overlap DEGs within three time points (A) and different two time points (B-D). Left panels show log_2_ (fold change), and right panels show FPKM values. Data are presented as mean ± SE from three biological replicates. Stars indicate significant differences between CK and HMF (calculated by DESeq2). *, *p* < 0.05; **, *p* < 0.01; ***, *p* < 0.001. The Araport11 annotations of functionally uncharacterized genes are noted below the corresponding pictures.

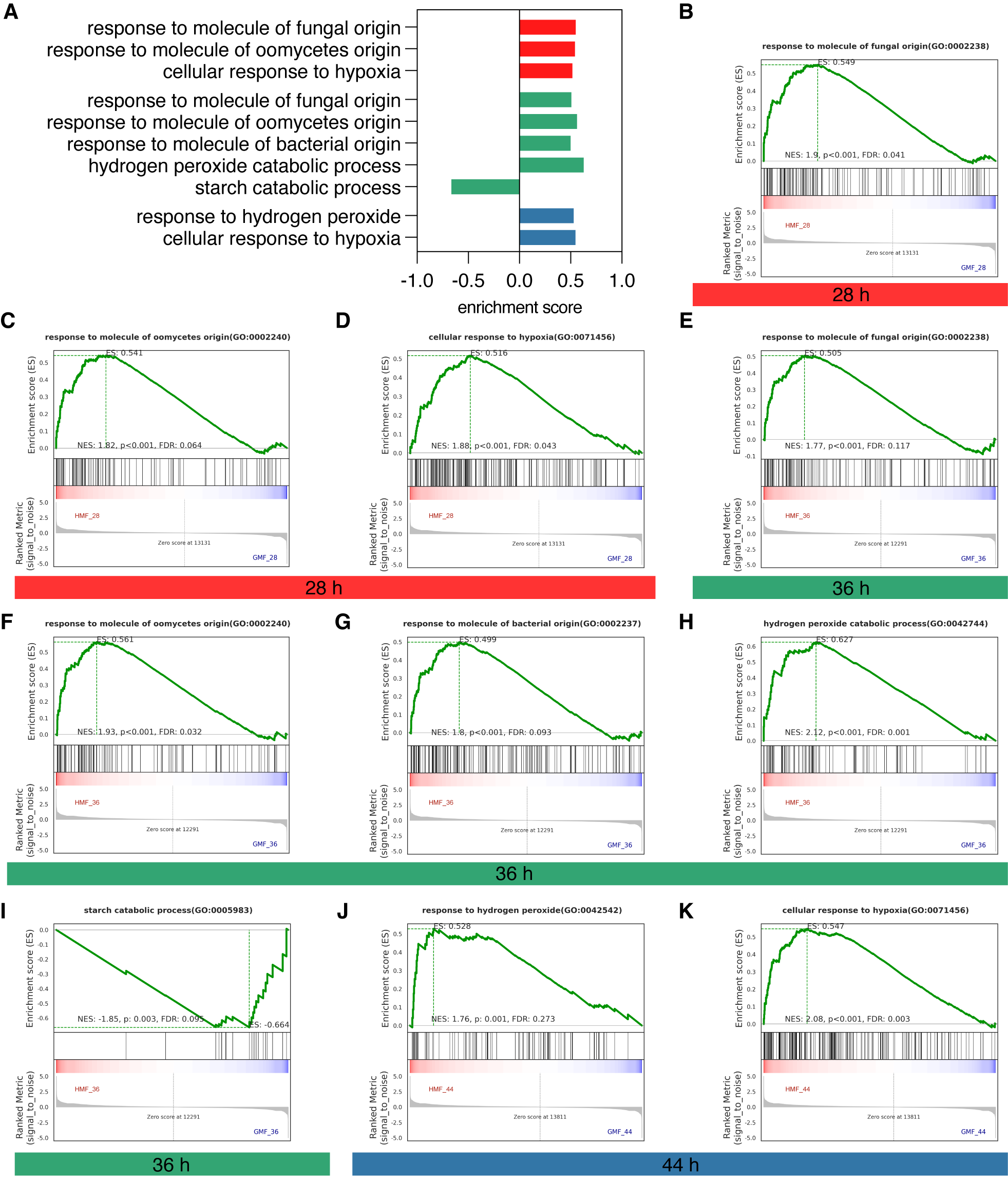

### Supplementary Fig. 10. GO process enriched in HMF by gene set enrichment analysis (GSEA).

(A) Summary of identified GO process significant enriched in HMF by GSEA. Red, green and blue bars represented 28 h, 36 h and 44 h sampling times. Positive enrichment scores indicate upregulation, while negative scores indicate downregulation in HMF.

(B-K) GSEA results of enriched process at 28 h (B, C), 36 h (D-I) and 44 h (J, K).

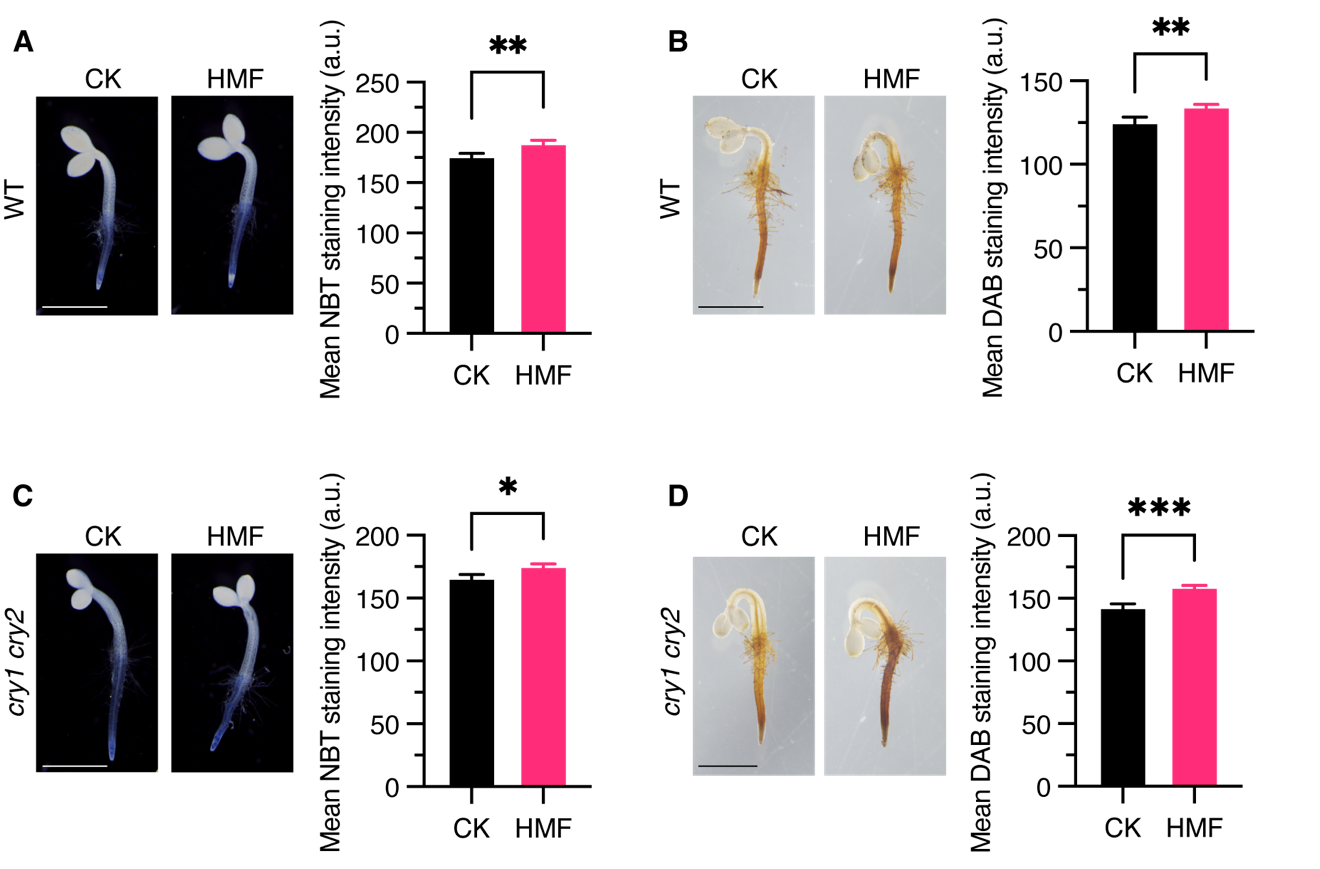

### Supplementary Fig. 11. *In situ* ROS staining of WT and *cry1 cry2* seedlings under white light.

(A, B) NBT (A) and DAB (B) staining and corresponding quantification data of WT seedlings grown in white light. Data are shown means ± SE (n ≥ 12). Stars indicate significant differences between CK and HMF (Student’s *t*-test). **, *p* < 0.01. Scale bar, 1 mm.

(C, D) NBT (A) and DAB (B) staining and corresponding quantification data of *cry1 cry2* seedlings grown in white light. Data are shown means ± SE (n ≥ 12). Stars indicate significant differences between CK and HMF (Student’s *t*-test). *, *p* < 0.05; ***, *p* < 0.001. Scale bar, 1 mm.

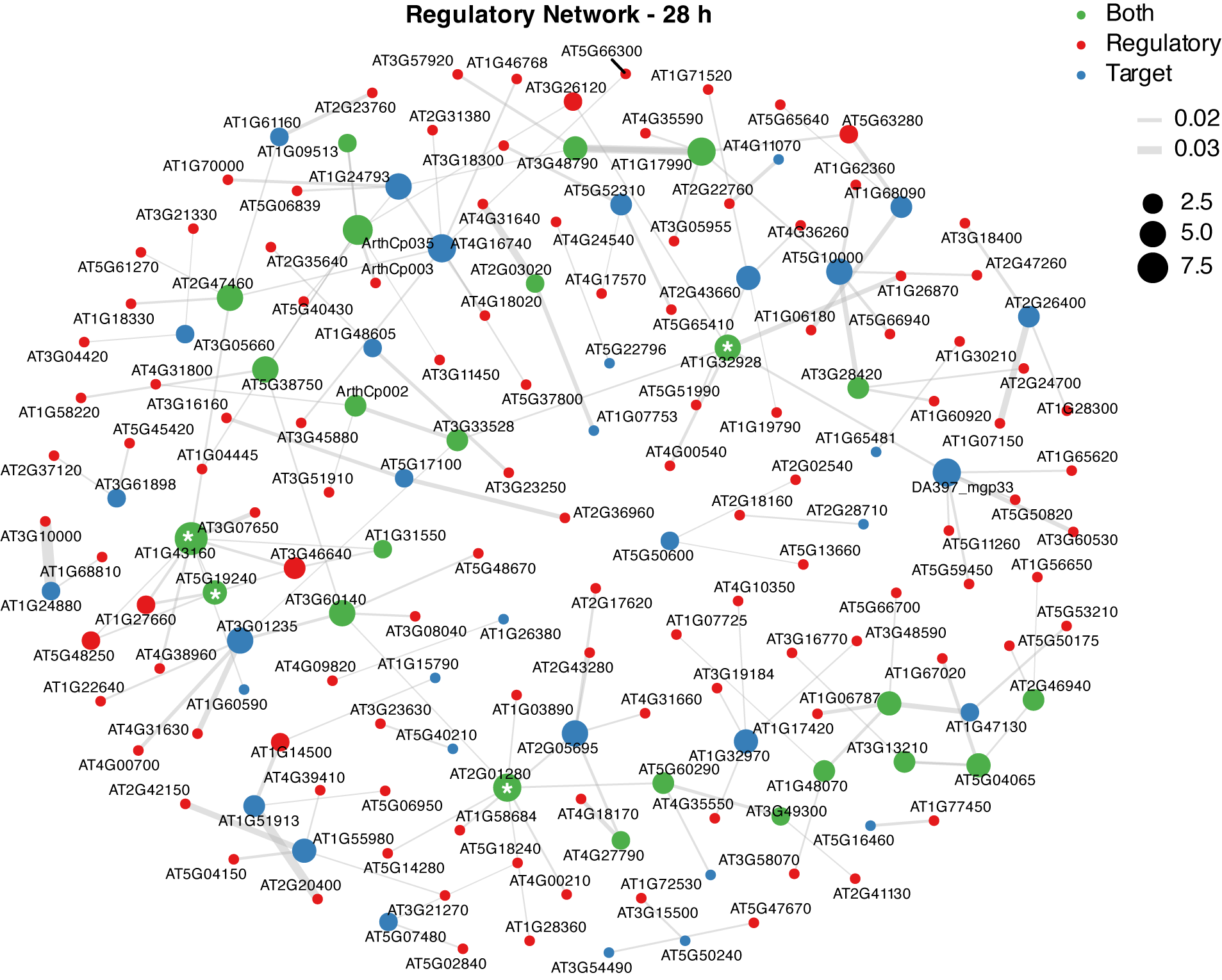

### Supplementary Fig. 12. HMF-specific regulatory network at 28 h.

Node color distinguishes regulators from targets; node size corresponds to the number of connected regulatory edges. Edge thickness indicates interaction strength. Stress-related regulators are marked with an asterisk (*).

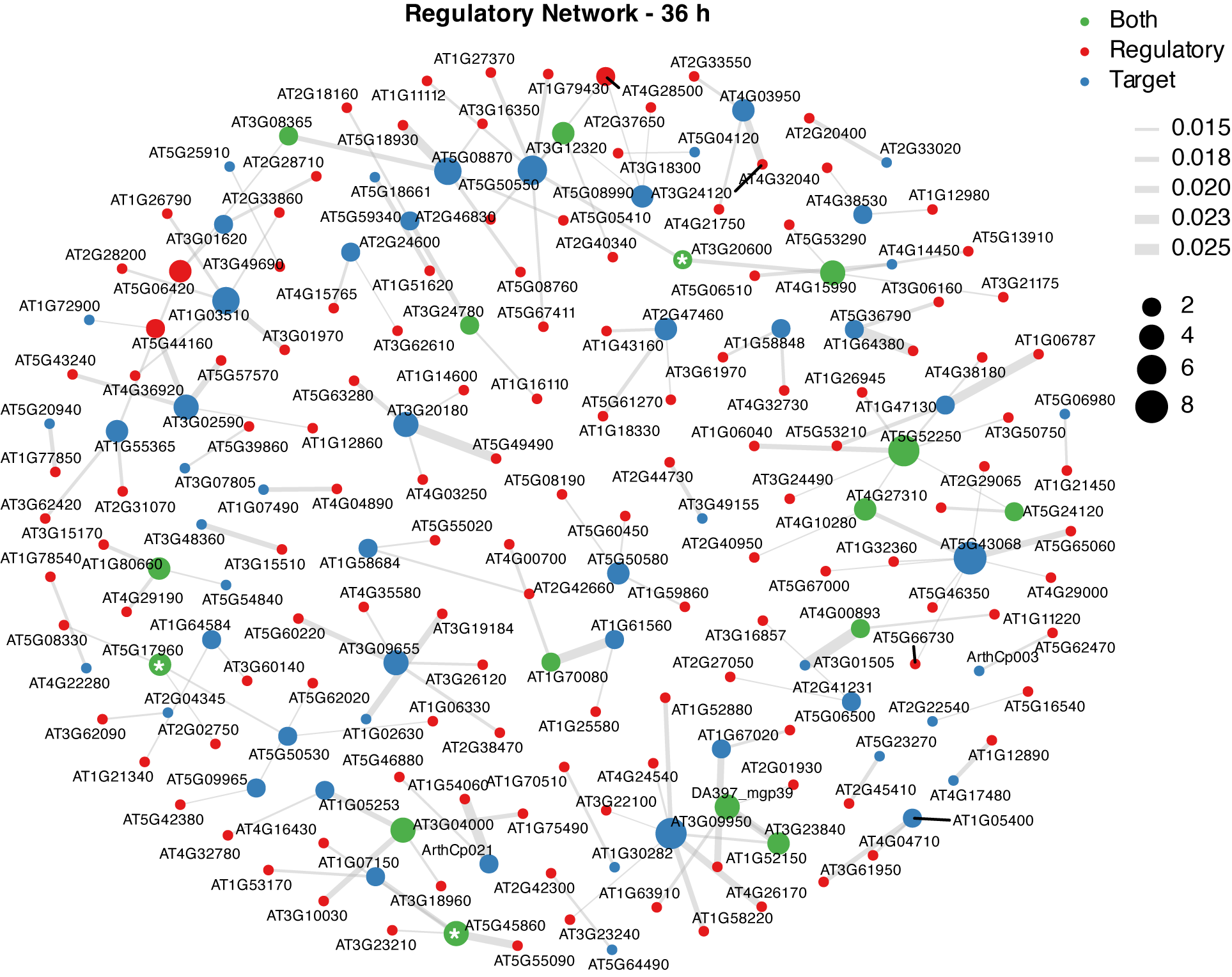

### Supplementary Fig. 13. HMF-specific regulatory network at 36 h.

Node color distinguishes regulators from targets; node size corresponds to the number of connected regulatory edges. Edge thickness indicates interaction strength. Stress-related regulators are marked with an asterisk (*).

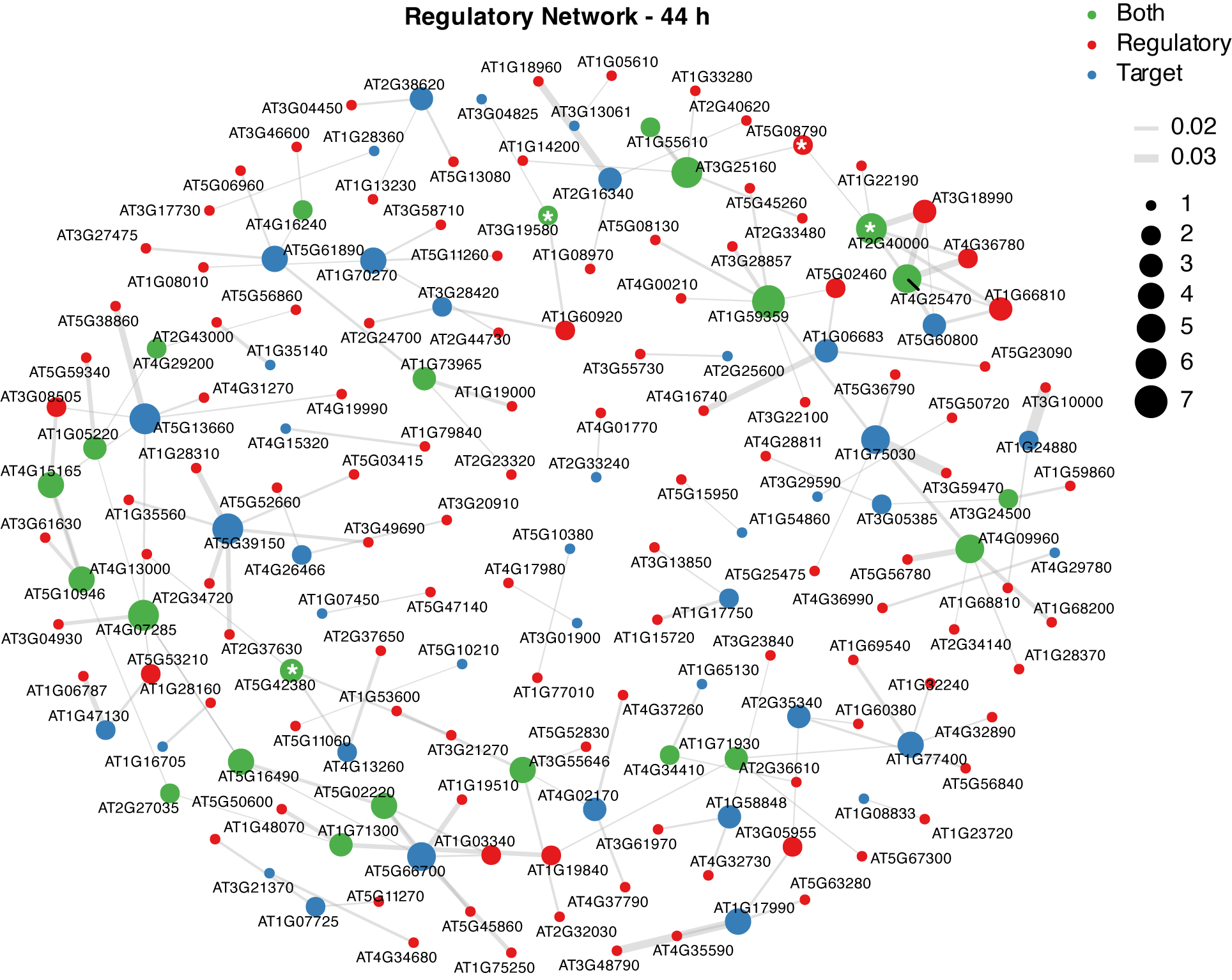

### Supplementary Fig. 14. HMF-specific regulatory network at 44 h.

Node color distinguishes regulators from targets; node size corresponds to the number of connected regulatory edges. Edge thickness indicates interaction strength. Stress-related regulators are marked with an asterisk (*).

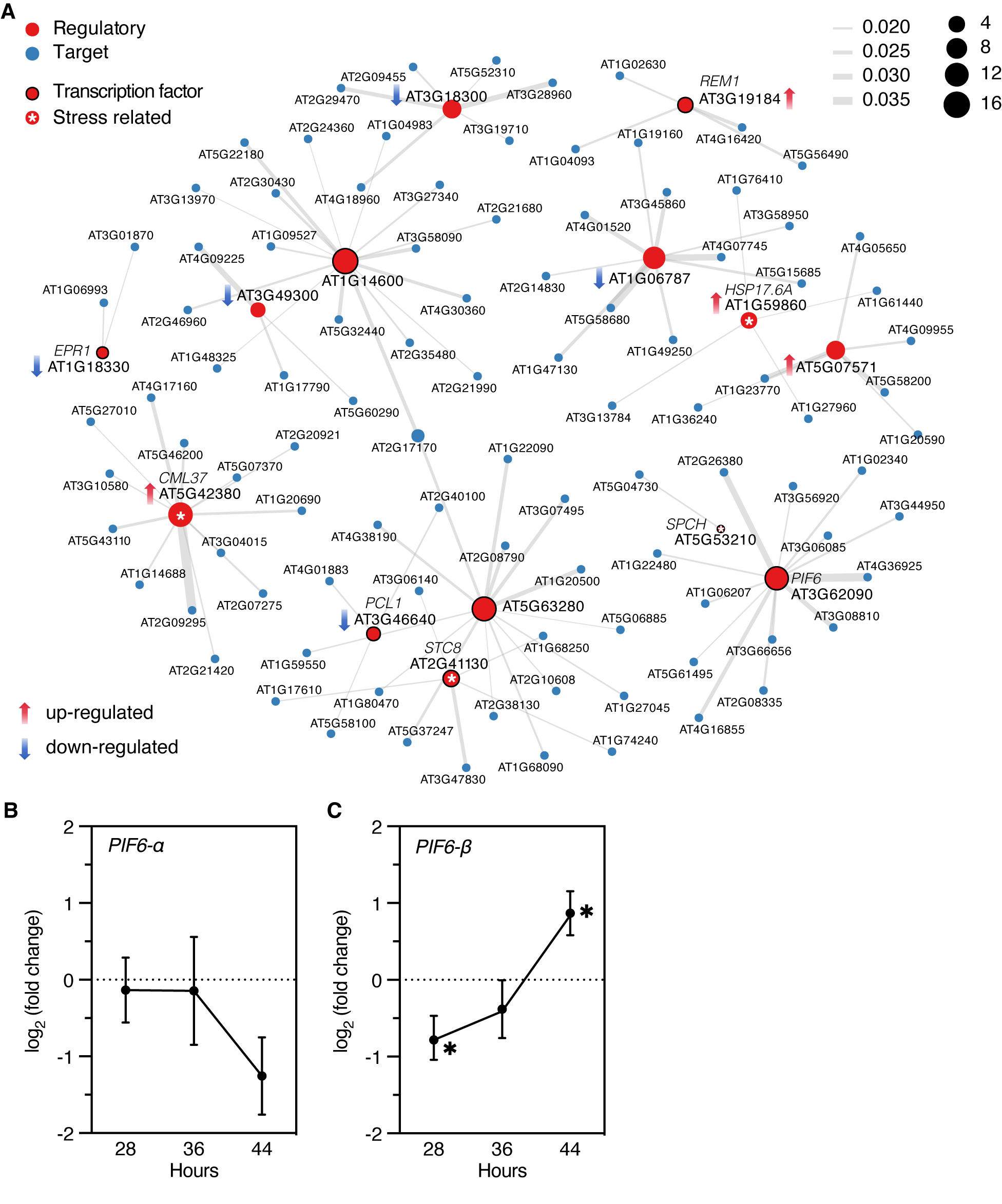

### Supplementary Fig. 15. Common regulator network and *PIF6* isoforms expression in HMF.

(A) HMF-specific regulatory network comprising regulators consistently activated across all three time points. Node color distinguishes regulators from targets; node size corresponds to the number of connected regulatory edges. Edge thickness indicates interaction strength. Stress-related regulators are marked with an asterisk (*). Transcription factors are outlined in black. Red and blue arrows indicating significant upregulation and downregulation under HMF conditions, respectively.

(B, C) Expression changes of *PIF6-α* (B) and *PIF6-β* (C) isoforms in HMF. Data are shown means ± lfcSE (n = 3). *, significant difference between CK and HMF (DESeq2, *p* < 0.05).

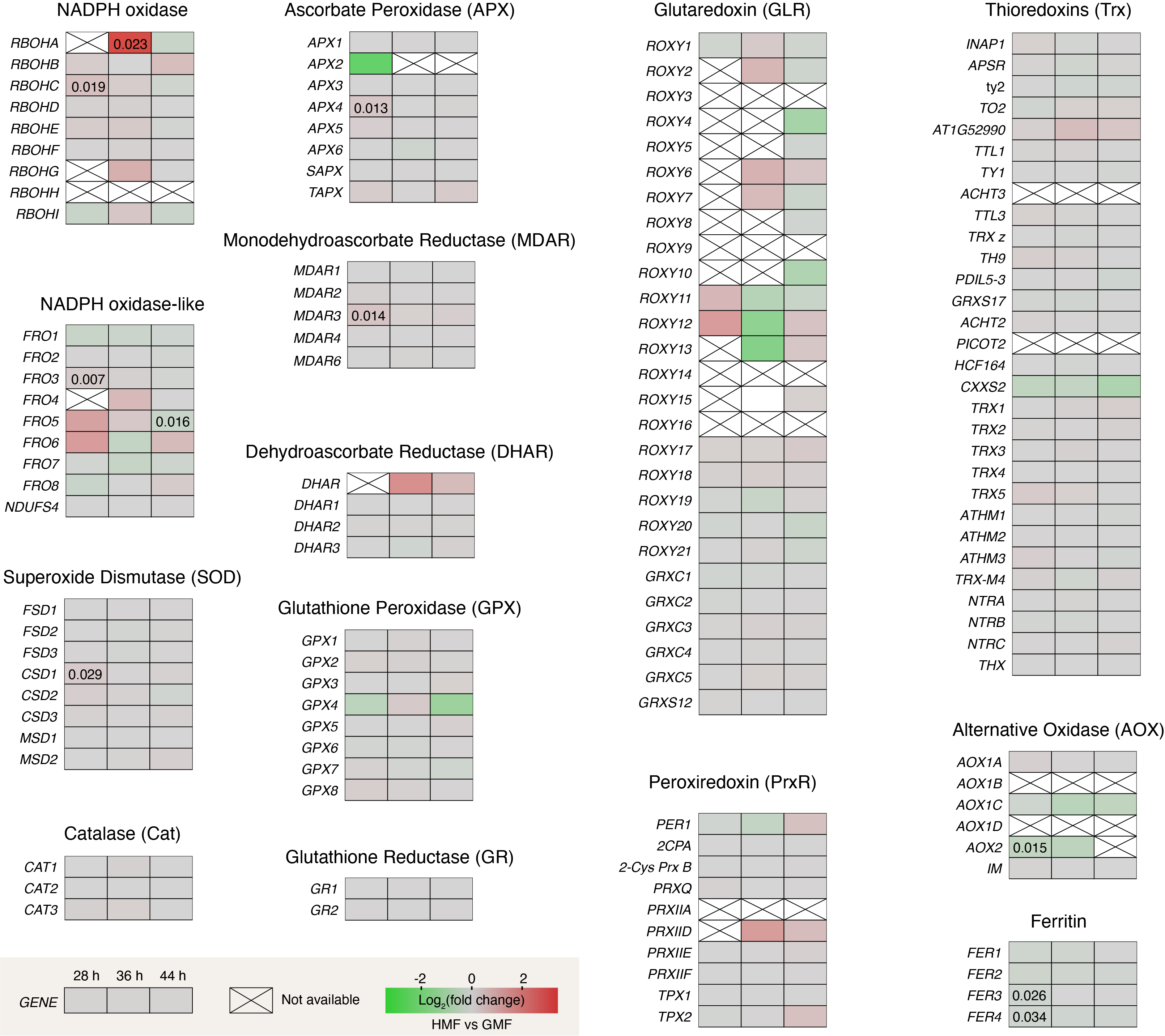

### Supplementary Fig. 16. Effects of HMF on the expression of enzymes involved in ROS metabolism.

Expression level changes (HMF vs GMF) for each gene at 28, 36 and 44 h, displayed from left to right in a panel. Red indicates upregulation while green means downregulation. The unavailable data are marked by a diagonal cross. *P*-value (DESeq2) less than 0.05 are labeled.
